# Time cells lead neural reinstatement of episodic memory

**DOI:** 10.64898/2026.06.28.734674

**Authors:** Aidan S. Dulaney, Sarah Seger, Oscar A. Carranza, Ruchita Mahesh Kumar, Joshua Jacobs, Bradley C. Lega

## Abstract

The episodic memory system provides humans with a unique ability to form and retrieve rich and detailed memories. This capacity requires representing temporal context at encoding and recovering it at retrieval to drive successful recall. Time cells and ramping cells in human medial temporal cortex have been proposed as substrates of temporal context, but whether they participate in context recovery during retrieval remains unknown. Using microelectrode recordings from neurosurgical patients across two classical episodic paradigms, free recall and serial reconstruction, we identify time sensitive neurons and demonstrate that these neuron populations participate in neuronal assemblies and organized sequences on gamma and theta time scales respectively. Consistent with recovery of temporal context predicted by behavioral models of episodic processing, time cell firing precedes the broader assembly population and initiates sequential activation retrieval. These properties predict contextually-mediated recall behavior. We also identify phase coding by complementary populations as convergent mechanisms whose recruitment depends on task demands. Finally, we demonstrate the emergence of context–sensitive neurons in a naturalistic viewing dataset lacking the predictable temporal scaffold of canonical assays. These findings identify cellular mechanisms by which the human medial temporal cortex represents temporal context during episodic encoding and recovers it during retrieval.

## Introduction

Episodic memory requires the brain to represent temporal context as events unfold and to recover that context at retrieval to drive successful recall. According to influential theoretical accounts, a shifting representation of temporal context binds to item information at encoding, and reinstatement of that context at retrieval cues the items associated with each contextual state ^1–3^. Subjects cluster their successive recalls by temporal proximity at encoding ^4,5^, and successful retrieval tracks reinstatement of the context present at encoding ^6,7^. It remains to be established how recovery of this context is implemented at the cellular level, and whether temporal context representation and recovery recruit different cellular mechanisms depending on task structure. To address these questions, we compared two episodic memory tasks, free recall and serial reconstruction, whose encoding periods are well-matched but differ in the functional role of temporal context. Free recall requires subjects to internally generate temporal context to cue item retrieval. Serial reconstruction provides item information and an external positional scaffold at retrieval, and the response order can in principle be supported by associative chaining between items without an internally generated temporal context ^8,9^.

Time cells and ramping cells have been proposed as cellular substrates of temporal context representation. Time cells fire at consistent moments within a structured time interval ^10^ and have been documented in rodent hippocampus and entorhinal cortex ^11–14^ and in human medial temporal cortex during episodic memory tasks ^15–17^. Ramping cells, also referred to as ‘temporal context cells’ in some published work, fire with monotonically changing rates over longer windows and have been identified across rodent, monkey, and human medial temporal and prefrontal cortex ^18–20^. Theoretical accounts cast these populations as components of a common temporal code, in which ramping cells implement a Laplace-domain representation of time and time cells emerge as the inverse-Laplace readout when the relevant interval is bounded and predictable ^18,19,21^.

A critical outstanding question is whether time cells and ramping cells participate in episodic retrieval by driving the recovery of temporal context. The temporal context model proposes that recovery of context at retrieval re-cues the items associated with that contextual state ^1–3^. At the cellular level, if reinstated context cues content within an integrative event, then context-carrying neurons should fire before content-carrying neurons within the integration window of a reactivating cell assembly. Cell assemblies are coordinated co-firing populations that have been proposed as integrative units of memory representation ^22–24^, with theta oscillations proposed as the temporal frame within which assembly integration occurs ^25–27^. Multi-neuron sequences extend the same integrative principle across longer timescales, with established roles in rodent memory ^12,28,29^ and emerging characterization in human medial temporal cortex ^30^. Prior work has shown that memory-sensitive neurons in human medial temporal cortex reinstate their encoding firing patterns at retrieval ^31^, consistent with the population-level reinstatement predicted by the temporal context model. However, it remains unknown whether this reinstatement involves an ordered within-cycle cellular signature– specifically whether context-carrying neurons firing before content-carrying neurons during individual reactivation events– and whether such ordering depends on task demands, has not been established. In free recall, where context must cue retrieval, this ordering should be essential. In serial reconstruction, where items are externally provided, context-driven ordering may be less critical.

While time cells and ramping cells represent temporal information through rate coding during encoding ^11,15^, an alternative account proposes that neurons may also carry temporal information through the phase at which they fire relative to ongoing theta oscillations ^25,32^. Initial intracranial reports identified phase relationships consistent with serial-position coding during working memory ^33^, but recent work has reframed these patterns as arising from interactions between stimulus timing and ongoing oscillations rather than reflecting an active ordering mechanism ^34^. Whether the medial temporal lobe recruits multiple temporal representations during encoding, and whether engagement of phase-based codes depends on task structure, remains unclear.

We tested these predictions by recording single units from human medial temporal cortex during two episodic memory tasks and a naturalistic viewing dataset. Specifically, we asked: (1) Do time cells and ramping cells carry rate-decodable temporal information, and do their properties differ between free recall and serial reconstruction? (2) Within reactivating cell assemblies, do time cells fire at earlier theta phases than other assembly members, and does this phase ordering depend on whether temporal context must cue retrieval? (3) Do multi-neuron sequences organize around time cells, and does their frequency track behavioral markers of contextually organized recall? (4) During encoding, do non-temporal neurons carry temporal information through theta-phase coding, and does the prevalence of phase-based representation depend on task structure?(5) Does classical temporal coding require bounded, predictable intervals, as theory predicts, or do they emerge across naturalistic temporal contexts?

## Results

### Time cells and ramping cells in human medial temporal cortex carry rate-decodable temporal information across both paradigms

We recorded single units from the medial temporal cortex of 62 human epilepsy patients performing one of two verbal memory paradigms (Figure 1A,B). In free recall (FR), participants studied lists of 12 or 15 words and verbally recalled them in any order during a 30–45 s retrieval period. In serial reconstruction (SR), participants studied lists of 10 words and reconstructed the encoding order from a randomized choice set. Our final samples comprised 768 units in free recall and 247 units in serial reconstruction. Behavioral performance followed canonical serial-position curves in both paradigms (Figure 1C), with better recall of early and late items in both paradigms, consistent with primacy and recency effects.

**Figure 1.**
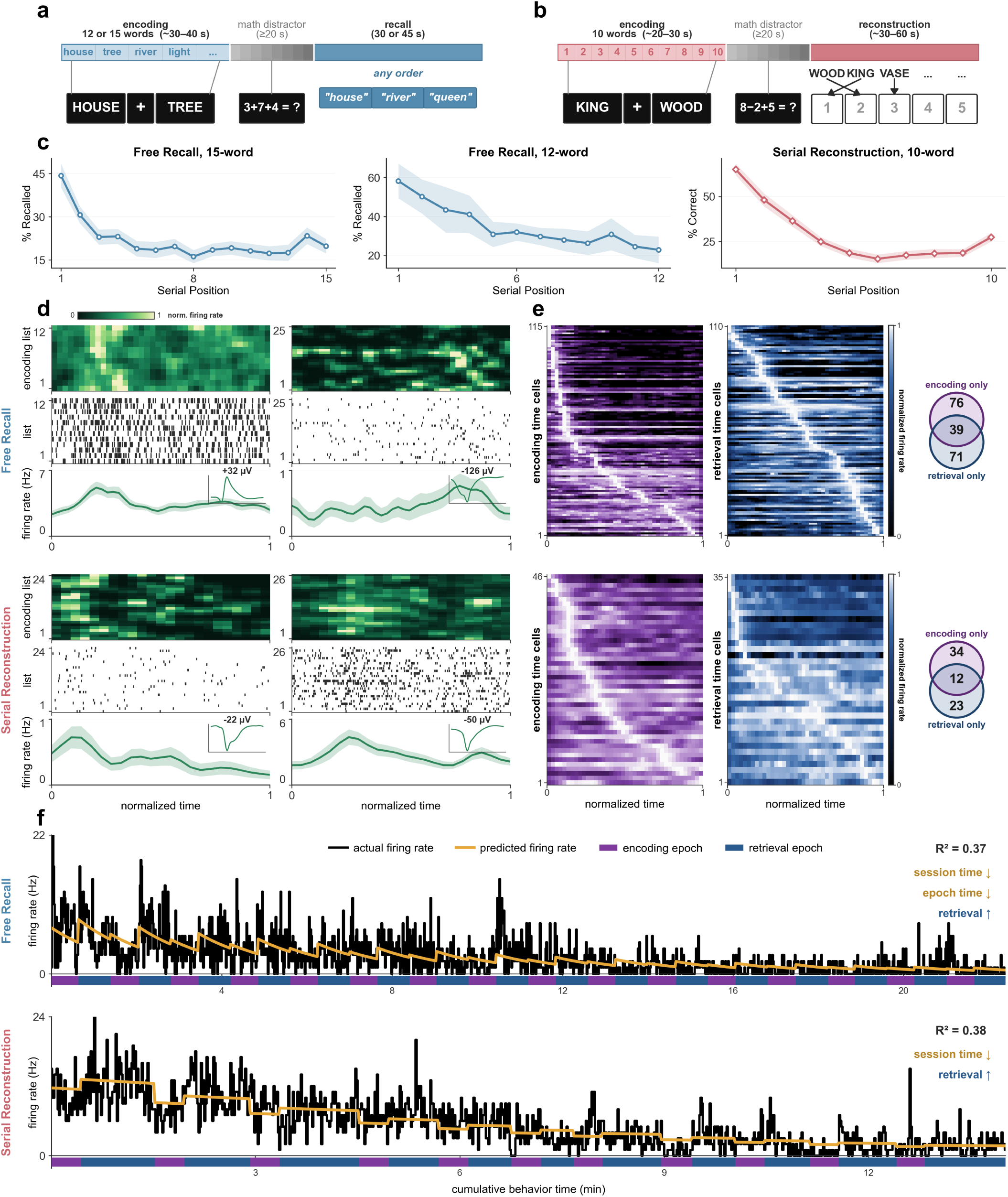
Behavioral paradigms and temporally tuned single units in human medial temporal cortex. (**a**) Free-recall task (FR): encoding of 12 or 15 words, a math distractor, then verbal recall in any order. (**b**) Serial-reconstruction task (FR): encoding of 10 words, a math distractor, then reconstruction of the studied order from a randomized choice set. (**c**) Serial-position curves for the 15-word and 12-word free-recall lists and the 10-word serial-reconstruction lists. (**d**) Example time cells in free recall (top) and serial reconstruction (bottom), showing per-list firing-rate heatmaps, spike rasters, trial-averaged firing rate, and the mean spike waveform (inset). (**e**) Normalized firing-rate heatmaps of the time-cell population tiling the encoding interval, sorted by peak time, separately for encoding (purple) and retrieval (blue); Venn diagrams give the overlap between encoding- and retrieval-active time cells. (**f**) Example ramping cells with observed (black) and GLM-predicted (orange) firing rates across the session and within encoding epochs.

We identified two classes of temporally tuned neurons: time cells, which fire at consistent latencies within encoding lists, and ramping cells, which monotonically modulate their firing rate across longer windows bounded by either epoch-time (list-time) or session-time. Both cell classes were identified using established criteria detailed in Methods. In free recall, 186 units met time-cell criteria (24.2% of 768), and in serial reconstruction 69 units did (27.9% of 247), with no significant difference in prevalence between paradigms (*χ*^2^ (1) = 1.18, p = 0.277). Time cells tiled the encoding interval in both paradigms (Figure 1D, E), with tuning peaks distributed across early, middle, and late portions of the interval. A significant fraction in each paradigm were active during both encoding and retrieval (20% in free recall, 18% in serial reconstruction, p< 0.01, binomial test), consistent with prior reports of context-dependent recruitment of partially overlapping time-cell subsets ^1,4,35^. Ramping cells in both paradigms exhibited monotonic firing-rate trajectories across the session and within encoding epochs (Figure 1F). Thus, both temporal-coding cell classes were recruited in both paradigms.

The temporal tuning properties of time cells and ramping cells differed systematically between paradigms. Time-field width as a fraction of the encoding interval was substantially smaller in free recall (median 10.6%, IQR 7.5–16.2%) than in serial reconstruction (median 47.0%, IQR 27.9–95.6%; Mann–Whitney *U* = 171, *p <* 1 *×* 10^−18^, Figure 2B), indicating that time-field width differs systematically with task. This task-dependent difference persisted when comparing absolute time-field widths (Supp. Figure 2A). Ramping-cell subtypes showed a complementary asymmetry between paradigms. Session-down ramps (cells whose firing rate declined across the session) were overrepresented in serial reconstruction (Figure 2C; *p <* 0.05), and epoch-down ramps (cells whose firing rate declined within each encoding epoch) were overrepresented in free recall (*p <* 0.01). Anterior-posterior differences in ramping-cell composition emerged, with posterior hippocampus showing higher overall ramping-cell prevalence and enrichment of epoch-down ramps, while anterior hippocampus showed relatively more session-down ramps; however, there was no difference in the ramping time constants (Supp. Figure 2B,C,E,F). Time-cell properties did not vary between anterior and posterior locations(Supp. Figure 2A,B,D). These results demonstrate that the two paradigms recruited differently tuned temporal-coding populations even where the cell-class definitions were identical, an asymmetry potentially attributable to the greater demand on internally generated within-list temporal context in free recall or the need to differentiate across item lists in multiple time scales.

**Figure 2.**
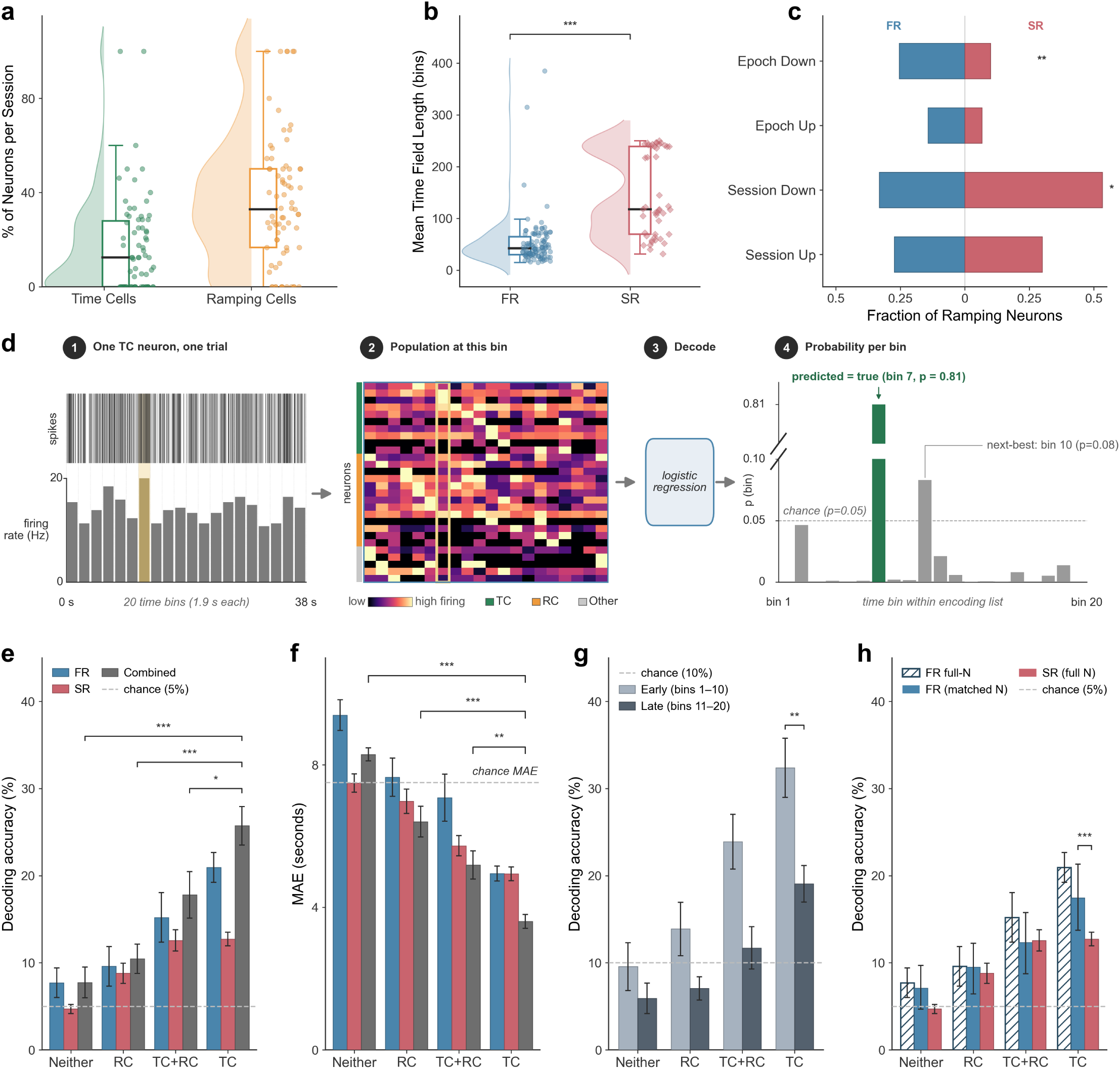
Tuning properties and population decoding of the encoding interval. (**a**) Proportion of recorded neurons per session classified as time cells and ramping cells. (**b**) Mean time-field width in free recall (FR) versus serial reconstruction (SR). (**c**) Distribution of ramping-cell subtypes (epoch/session, up/down) across paradigms. (**d**) Schematic of the population time-bin decoder: per-trial population firing rates are passed to a logistic-regression classifier that predicts the within-list time bin. (**e**) Decoding accuracy and (**f**) mean absolute error for four cell pools (neither, ramping cells, time + ramping, time cells) in FR, SR, and the combined sample; dashed lines indicate chance. (**g**) Decoding accuracy for early versus late list bins. (**h**) Decoding accuracy after subsampling the FR time-cell pool to match SR population size.

We trained a logistic-regression decoder to predict the within-list time bin (20 bins) from population firing rates and applied it separately to four cell-type pools (Figure 2D). In both paradigms, the time-cell pool decoded the encoding interval more accurately than ramping cells alone, the union of the two, or a control pool of cells classified as neither (Figure 2E,F). The accuracy hierarchy held for both decoding accuracy and mean absolute error, indicating that the rate-decodable temporal signal is concentrated in time cells rather than distributed across all temporally tuned populations. Decoding accuracy was higher for early than for late list positions (Figure 2G; ^∗∗∗^*p <* 0.001), echoing the behavioral primacy effect ^4,5^ and indicating that decodable temporal information is non-uniformly distributed across the encoding interval. The narrower time fields in free recall (Figure 2B) predicted higher per-cell decoding accuracy in this paradigm relative to serial reconstruction. To test this prediction independent of population size, we randomly subsampled the free-recall time-cell pool to match the number of serial-reconstruction time cells and reran the decoder. The per-cell decoding accuracy remained significantly higher in free recall compared to serial reconstruction (2H; ^∗∗∗^*p <* 0.001), consistent with free recall’s narrower time-field widths.

### Time cells phase-precede assembly reactivation during episodic retrieval

The temporal context model proposes that successful retrieval depends on the initial recovery of the contextual state present at encoding, which in turn re-cues the items ^1–3,6,7^. Cell assemblies—transient co-firing populations that recur across encoding and retrieval—are integrative units through which both item and context information are jointly expressed ^22–24^. Assembly activation is organized by theta oscillations, which govern when assemblies activate ^36^. If temporal context recovery initiates item reinstatement at the cellular level, then context-carrying neurons should fire at an earlier theta phase than the broader assembly population. To test this prediction, we identified assemblies (3A) and examined whether time cells fire at a systematically earlier theta phase than other assembly members during retrieval reactivation events. Following established methods, we identified cell assemblies as groups of neurons co-firing within 25 ms bins during encoding more than expected for independently firing cells ^36,37^. We defined reactivation events as moments during retrieval when assembly-weighted population activity exceeded a per-assembly threshold ^38^. Across sessions, these assemblies each comprised a stable fraction of the recorded population, and assembly members, but not nonmembers, tracked the strength of reactivation (Supp.F̃igureS̃ 3A,B).

For each cell assembly, we identified the preferred theta frequency as the frequency within the theta band at which its member neurons were most strongly phase-locked to the local field potential. For each reactivation event of that assembly, we then computed the phase difference between time-cell spikes within a 150 ms window of the onset of reactivation and the population mean phase of the assembly’s contributing neurons, with negative values indicating time-cell spikes occur at an earlier phase. Time cells fired at a systematically earlier theta phase than the surrounding assembly population in both paradigms (Figure 3B). In free recall, the mean shift was Δ*θ* = *−*63^◦^, approximately *−*29 ms at 6 Hz (MRL = 0.18, cluster-bootstrap *p* = 0.001, 80 assemblies across 19 subjects). In serial reconstruction, the mean shift was Δ*θ* = *−*23^◦^, approximately *−*11 ms (MRL = 0.48, cluster *p* = 0.002, 42 assemblies across 12 subjects). The magnitude of the phase lead was larger in free recall than in serial reconstruction (Δ*θ* = *−*63^◦^ vs. Δ*θ* = *−*23^◦^) but was present in both, indicating that time-cell phase precedence is a feature of assembly reactivation independent of whether temporal context is internally generated. This lead did not depend on assuming an oscillation or on specific analysis parameters: it was recovered when measured directly in spike time (*≈ −*41 ms in free recall, *≈ −*9 ms in serial reconstruction), held across fixed theta frequencies from 3 to 9 Hz and across activation-window widths, and was present on the majority of individual reactivation events of a given assembly (Supp.F̃igureS̃ 3C–F,H). In contrast, ramping cells revealed a smaller phase lead in serial reconstruction (Δ*θ* = *−*10^◦^, approximately *−*5 ms, cluster *p* = 0.005) but no consistent phase relationship in free recall. The temporal organization of time cells, but not ramping cells, around assembly reactivation is consistent with the cellular prediction of the temporal context model in both paradigms, raising the mechanistic question of whether this organization shapes how strongly the assembly itself reactivates.

**Figure 3.**
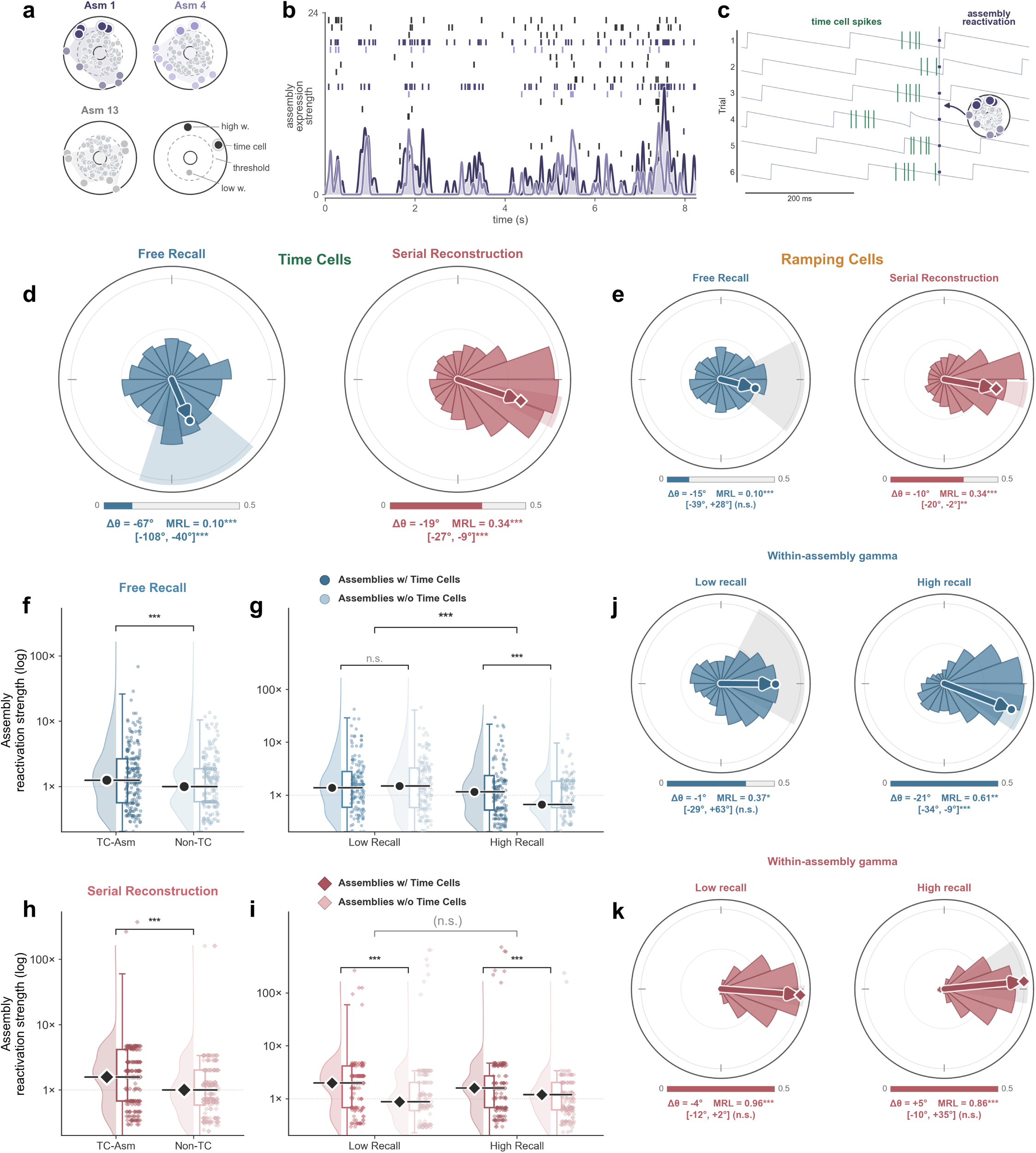
Time cells phase-lead assembly reactivation during retrieval. (**a**) Assembly detection (PCA–ICA): example assemblies with member weights, threshold, and time-cell membership. (**b**) Assembly expression strength over a retrieval period. (**c**) Gamma-band spiking around a reactivation event. (**d**) Theta phase of time-cell spikes relative to the assembly population mean for FR and SR; negative Δ*θ* indicates a phase lead. (**e**) The same comparison for ramping cells. (**f**, **h**) Assembly reactivation strength for assemblies containing at least one time cell (TC-Asm) versus none (Non-TC), in FR (**f**) and SR (**h**). (**g**, **i**) The same comparison split by low- and high-recall lists, in FR (**g**) and SR (**i**). (**j**, **k**) Within-assembly slow-gamma phase of time-cell versus non-time-cell members on low- and high-recall lists, in FR (**j**) and SR (**k**).

Previous work has shown that the strength of assembly reactivation at retrieval tracks behavioral memory expression ^35,38^. If time cells provide the contextual cue for assembly reactivation, assemblies containing time cells (TC-Asm) should reactivate more strongly than those without (non-TC assemblies), with this advantage scaling with successful recall. We compared reactivation strength (projection of instantaneous population activity onto assembly weight vector) between TC-Asm and non-TC assemblies, dividing lists by median recall performance. TC-Asm reactivated more strongly than non-TC assemblies in both paradigms (Mann–Whitney *U*, *p <* 0.001 in both; Figure 3C,D). This advantage was not attributable to assembly size, which was comparable for time-cell-containing and other assemblies (Supp.F̃igureS̃ 3G). A linear mixed-effects model with subjects as a random effect tested whether this TC-versus-non-TC advantage was modulated by recall success. In free recall, the interaction was significant (LMM *p <* 0.001). The TC-Asm advantage was 1.80*×* larger on high-recall than on low-recall lists, with the two assembly types reactivating indistinguishably on low-recall lists and TC-Asm reactivating 1.70*×* more strongly than non-TC assemblies on high-recall lists (Figure 3C). In serial reconstruction, the TC advantage was present but independent of recall (LMM p = 0.69), consistent with weaker contextual dependence when items are externally provided (Figure 4C,D).

**Figure 4.**
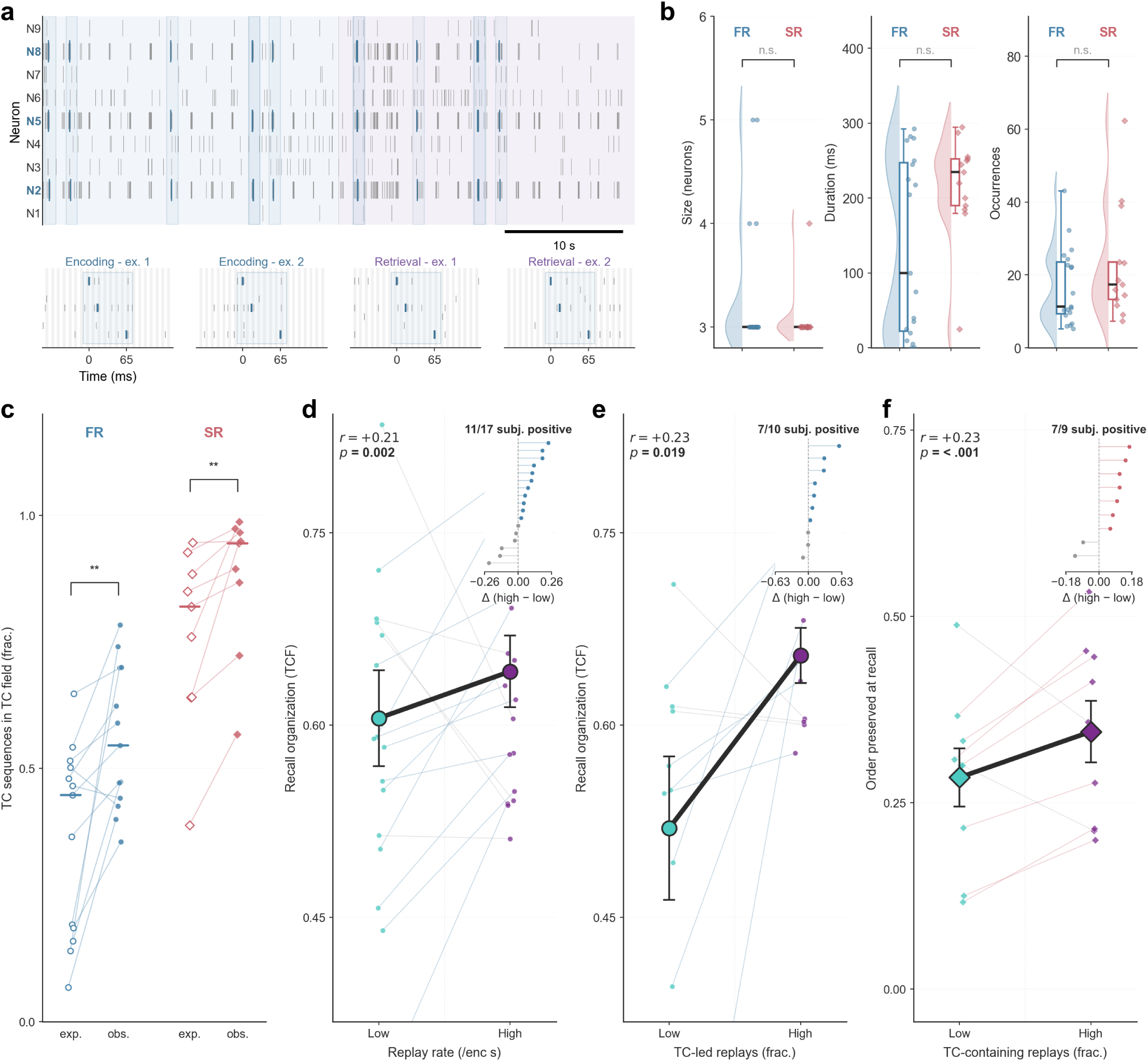
Time-cell-led multi-neuron sequences track contextually organized recall. (**a**) Example detected sequence across neurons, recurring across encoding and retrieval, with expanded views of individual sequence occurrences. (**b**) Sequence size, duration, and occurrence count in FR versus SR (n.s.). (**c**) Fraction of sequences engaging time cells within their time fields, observed versus expected. (**d**) Within-subject relationship between replay rate and temporal clustering factor (TCF). (**e**) TCF as a function of the fraction of time-cell-led replays. (**f**) Order accuracy as a function of the fraction of time-cell-containing replays in SR. Insets show per-subject high-minus-low differences.

These analyses demonstrated that time cells precede assembly reactivation at retrieval and that TC-containing assemblies predict free-recall success. We then looked within the assemblies themselves, positing that time cells should fire earlier within a cycle defined by low gamma oscillations. Gamma oscillations define the integration window within which assembly members are bound ^23,25,39^, so we compared the gamma phase at which time-cell members fired to that of their non-time-cell co-members within each reactivating assembly. Phase was estimated at 40 Hz, the center frequency used for within-assembly phase comparisons in human medial temporal cortex ^36^. In most conditions, time-cell and non-time-cell members fired at indistinguishable gamma phases (Δ*θ* near 0^◦^, cluster *p >* 0.1 for low performing lists in free recall (Figure 3E, left) and for low and high performing lists in serial reconstruction (Figure 3F). The exception was free recall on high-recall lists, where time cells led their non-time-cell co-members by Δ*θ* = *−*21^◦^, approximately *−*1.5 ms (MRL = 0.61, cluster *p <* 0.001, 16 assemblies across 7 subjects; Figure 3E). The gamma-scale precedence thus appears only during successful free-recall retrieval, consistent with a finer-timescale extension of the theta-scale precedence documented above.

### Time cells lead multi-neuron sequences whose abundance tracks behavioral measures of contextual recall

We examined whether multi-neuron sequences initiated by time cells predict contextually-organized recall behavior ^30^. This tests whether the same time-cell-leading principle that organizes individual assemblies extends to longer timescale retrieval chains that structure behavioral memory. We applied the Spike Pattern Detection and Evaluation (SPADE) algorithm to detect repeating temporal spike patterns against a dithered surrogate null (Methods) ^40,41^. Significant sequences were detected in both paradigms, with the same patterns recurring across both encoding and retrieval epochs (Figure 4A). Pattern size, duration, and per-pattern occurrence count did not differ between paradigms (Figure 4B; free recall, 3.4 *±* 0.7 neurons, 125 *±* 82 ms, 17 *±* 10 occurrences per pattern; serial reconstruction, 3.1 *±* 0.2 neurons, 185 *±* 42 ms, 23 *±* 16 occurrences; all *p >* 0.05, Mann–Whitney *U*).

We next asked whether the neurons participating in sequences were drawn preferentially from the assemblies identified above. These neurons were significantly more likely to be assembly members than the recorded population at large (Supp. Figure S4A; ^∗∗∗^*p <* 0.001 in both paradigms), and within sequences they were significantly more likely to be drawn from the same assembly than from different assemblies (Supp. Figure S4B; observed *>* shuffled, ^∗∗∗^*p <* 0.001 in both). Time cells participated in sequences preferentially within their time fields, consistent with their tuning to specific moments of the encoding interval (Figure 4C; Wilcoxon signed-rank, ^∗∗^*p <* 0.01 in both paradigms). A sequence that recurred during both encoding and retrieval of the same list, defined as a replay event, occurred more often than expected under a dither-surrogate null in both paradigms (Supp.F̃igureS̃ 4C,D; free recall $1.31\times$, serial reconstruction $1.12\times$). The frequency of these replay events across encoding and retrieval is a candidate cellular index of contextually organized recall. We tested whether it tracks each subject’s temporal clustering factor (TCF), a behavioral measure of how consistently successive recalls cluster around nearby encoding positions and a hallmark behavioral signature of retrieved-context retrieval ^4,5^. In free recall, the within-subject correlation between replay frequency and TCF was positive (Figure 4D; cluster-bootstrapped *r* = +0.21, ^∗∗^*p* = 0.002, 11 of 17 subjects positive).

If time cells are central to the contextual organization that replay supports, restricting to replays *initiated by time cells* should preserve or strengthen the behavioral correlation. We defined such time-cell-led replays as sequences whose first participating neuron is a time cell. Within this subset, the correlation with TCF was preserved (Figure 4E; *r* = +0.23, ^∗^*p* = 0.019, 7 of 10 subjects positive). Within each subject, splitting lists by replay rate revealed a larger TCF difference between high- and low-replay halves under the time-cell-led predictor than under the broad replay rate (paired Wilcoxon signed-rank across *n* = 10 subjects, median ΔTCF_TC-led_ *−* ΔTCF_broad_ = +0.08, mean +0.11, 95% bootstrap CI [+0.03, +0.21], *p* = 0.028, 8 of 10 subjects positive), confirming that the time-cell-led subset carries the contextual-organization signal more strongly than the broader replay rate. In serial reconstruction, the analogous correlation between time-cell-containing replay fraction and behavioral order accuracy was positive (Figure 4F; *r* = +0.23, ^∗∗∗^*p <* 0.001, 7 of 9 subjects positive).

### Theta-phase coding represents temporal context across serial positions and within discovered task structure

Theta oscillations provide a complementary mechanism through which temporal information may be organized during encoding. Theoretical accounts propose that the theta phase at which a neuron spikes may encode sequential position, with successive items represented at progressively shifted phases within each theta cycle ^25,42^, and MTL neurons reliably phase-lock to theta during memory encoding ^31,43^. Whether phase organizes temporal or sequential information in humans, however, remains unclear. Because time-sensitive neurons exhibit phase precession. a distinct phase-related phenomenon, we restricted this analysis to non-temporal neurons (i.e., neurons classified as neither time cells nor ramping cells). We tested for: (1) consistent phase differences across serial positions using phase locking analysis, and (2) graded relationships between spike phase and serial position using circular-linear correlation (Figure 5A, B).

**Figure 5.**
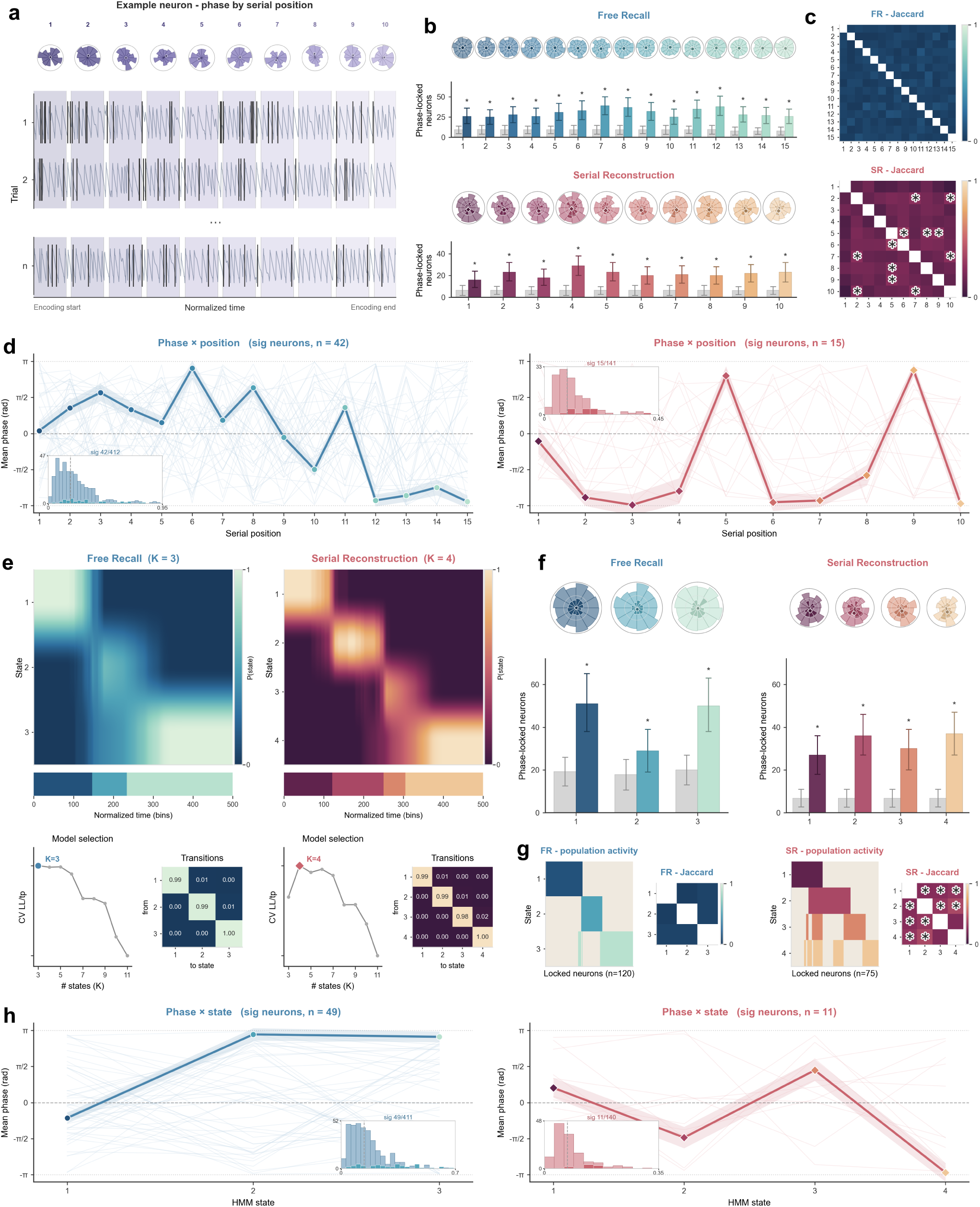
Theta-phase coding across serial positions and HMM-defined population states. (**a**) Example non-temporal neuron: spike phase relative to theta across serial positions, with single-trial rasters. (**b**) Number of significantly phase-locked neurons at each serial position in free recall (top) and serial reconstruction (bottom); grey bars indicate the shuffle null. (**c**) Pairwise Jaccard overlap of phase-locked populations across serial positions (asterisks, FDR-corrected *p <* 0.05). (**d**) Mean preferred phase as a function of serial position for significantly correlated neurons; insets show the distribution of circular–linear correlation values. (**e**) Hidden Markov models of population activity (FR, *K* = 3; SR, *K* = 4), with state-probability maps, cross-validated model selection, and transition matrices. (**f**) Phase-locked neuron counts per HMM state in free recall and serial reconstruction. (**g**) Population-activity and Jaccard summaries across HMM states. (**h**) Mean preferred phase as a function of HMM state.

In serial reconstruction, the number of cells showing significant phase non-uniformity exceeded chance at all serial positions (*p_perm_ <* 0.05; Figure 5B), whereas in free recall, this occurred only at positions 5–9 and 11–15. The proportion of neurons showing significant theta phase-locking was significantly higher in serial reconstruction (21/141, 14.9%) than in free recall (28/413, 6.8%; *χ*^2^(1) = 8.58, p = 0.003). However, the mean preferred phase did not vary systematically across serial positions in either paradigm, suggesting neurons do not encode position through phase shifts (serial reconstruction: grand MRL of state mean vectors = 0.859, *p_perm_* = 0.287; free recall: grand MRL of state mean vectors = 0.061, *p_perm_* = 0.262; Figure 5B). Pairwise comparisons of phase-locked populations across serial positions (Jaccard similarity) revealed no significant overlap in free recall after FDR correction (all FDR corrected p > 0.05). In serial reconstruction, six position pairs showed significantly above-chance overlap (Figure 5C; FDR-corrected Jaccard similarity, p < 0.05), concentrated around positions 2, 5, 7, and 10. Together, these results indicate that phase-locked populations are largely position-specific in free recall, but that serial reconstruction additionally recruits a subset of shared neuronal populations across select serial positions.

We tested whether phase carries graded positional information within individual lists using circular-linear correlation between theta phase and serial position. A small but significant proportion of neurons showed this relationship in both paradigms (serial reconstruction: 10.6% sig (15/141), *p_perm_* = 0.005; free recall: 10.2% sig (42/413), *p_perm_ <* 0.001; Figure 5D), with no significant difference between conditions (10.6% vs. 10.2%; *χ*^2^(1) = 0.025, p = 0.874). This suggests both paradigms recruit phase-based positional coding at the single-neuron level.

Serial position is an experimenter-defined variable that may not match how subjects naturally segment the encoding period. A phase code organized around internally defined population states would be obscured when parsed by serial position. We therefore tested whether theta phase coding is organized by latent population states rather than serial position. To identify these states, we fit an unsupervised hidden Markov model (HMM) to the full recorded population activity, including time cells and ramping cells. Model order was determined by cross-validated log-likelihood, which favored three states in free recall and four in serial reconstruction (Figure 5E). State-probability maps revealed that each state dominated a distinct contiguous portion of the encoding interval, tiling the period sequentially, and transition matrices confirmed the feed-forward structure of the model (Figure 5E).

We repeated the phase locking and circular-linear correlation analyses described above, substituting HMM state identity for serial position. In serial reconstruction, neurons showed significant phase locking at all HMM states (all FDR corrected *p_perm_ <* 0.05, Figure 5F), with no significant difference in mean preferred phase across states (grand MRL of state mean vectors = 0.962, *p_perm_* = 0.144; Figure 5F). Phase-locked populations showed significant overlap between most state pairs (FDR-corrected p < 0.05, Figure 5F) except states 3–4 (FDR-corrected p > 0.05). In free recall, significant phase locking occurred only at states 1 and 3 (FDR-corrected p < 0.05, Figure 5F), with no systematic phase shift across these states (grand MRL of state mean vectors = 0.272, *p_perm_* = 0.096; Figure 5F). Phase-locked populations showed no significant overlap between any state pairs in free recall (all FDR corrected p > 0.05; Figure 5F). The proportion of neurons showing significant theta phase-locking was significantly higher in serial reconstruction (33/141, 23.4%) than in free recall (50/413, 12.1%; *χ*^2^(1) = 10.53, p = 0.001). For circular-linear correlation between theta phase and HMM state, a small proportion of neurons showed significant relationships in both paradigms (serial reconstruction: 7.8% sig (11/141), *p_perm_* = 0.115; free recall: 11.9% sig (49/413), *p_perm_ <* 0.001; Figure 5E), with no significant difference between conditions (7.8% vs. 11.9%; *χ*^2^(1) = 1.80, p = 0.180). Overall, our interpretation is that phase encoding of serial position emerges in reconstruction, whereas in free recall phase coding develops more so from latent states defined by internal context generation.

### Naturalistic viewing recruits ramping cells

The cellular signatures of temporal context representation and recovery characterized above were obtained in paradigms with bounded, repeatable encoding intervals, the kind of temporal scaffold against which classical time-cell tuning is conventionally defined ^11,12^. We next asked what happens to the temporal-coding architecture when this scaffold is absent, by examining a separate dataset in which patients viewed a 28-minute episode of *Curb Your Enthusiasm* (CURB; 90 units across 12 patients; Methods). The episode comprised 17 scenes of unequal duration without repeated structure across the recording (Figure 6A), and patients made temporal judgments about the episode rather than encoding a structured list. This paradigm removed the bounded, repeatable interval that conventionally supports time-cell tuning while preserving a temporal demand on the participant.

**Figure 6.**
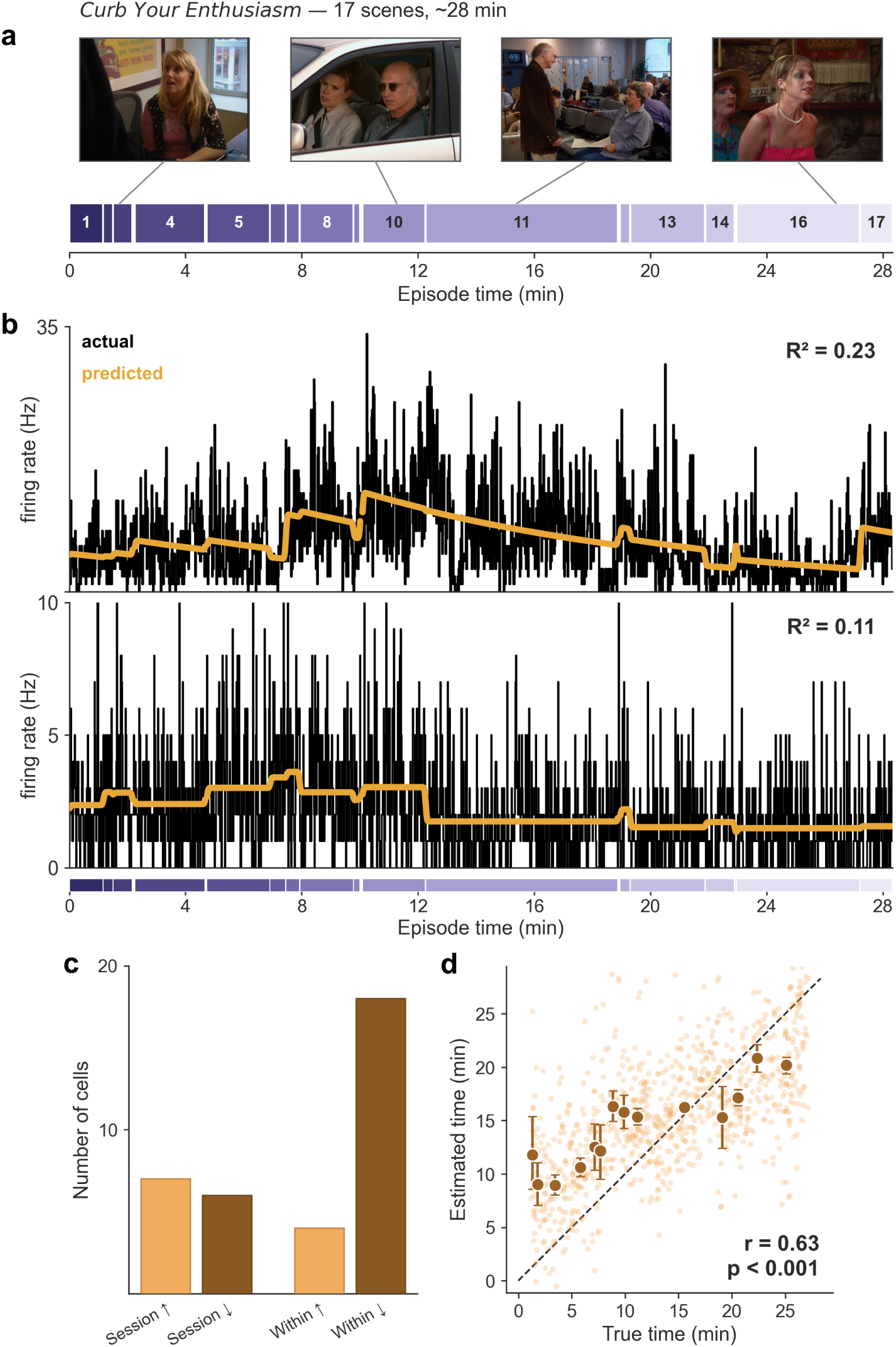
Naturalistic viewing recruits ramping cells but not classical time cells. (**a**) Scene structure of the 28-minute *Curb Your Enthusiasm* episode (17 scenes of unequal duration). (**b**) Example ramping cells with observed (black) and GLM-predicted (orange) firing rates: a session-scale ramp (top) and a within-scene ramp (bottom). (**c**) Counts of ramping-cell subtypes by timescale (session vs within-scene) and direction (up vs down). (**d**) Decoded versus true elapsed time across the episode.

We focused on ramping cells, which may provide a more suitable framework for examining temporal representation across naturalistic timescales. Notably, scene boundaries in the episode provided natural structural breaks analogous to list boundaries in the episodic tasks. We therefore modeled ramping-cell firing using the same stepwise Poisson GLM extended to include scene-onset and within-scene-time predictors. Ramping cells were identified in 32 of 90 units (35.6%) using the same stepwise Poisson GLM extended to include scene-onset and within-scene-time predictors (Methods; Figure 6B). The proportion of ramping cells identified in this paradigm was similar to that of free recall and serial reconstruction (Supp. Figure 5A).

The detected ramping population was heterogeneous across timescales. Thirteen cells (14.4%) showed session-scale ramps spanning the full 28-minute episode, and 22 cells (24.4%) showed within-scene ramps reset at scene onsets. Both upward and downward ramps were detected within each timescale, with within-scene downward ramps the most prevalent subtype (18 cells, 20.0%; Figure 6B). The functional distinction between upward and downward ramping cells in human medial temporal cortex remains poorly characterized, as down ramping neurons are mostly considered in models of temporal context representation built from these populations. Single-cell examples illustrated both organizational scales, including a session-scale upward ramp (Figure 6D, top; *p* = 1.3 *×* 10^−6^) and a within-scene downward ramp (Figure 6D, bottom; *p* = 7.6 *×* 10^−17^). These observations suggest that ramping-cell activity provides a continuous temporal representation that operates flexibly across multiple timescales, persisting even when explicit task structure and bounded intervals are absent.

## Discussion

How the human brain represents the temporal structure of experience at the cellular level, and how it recovers that structure to guide retrieval, has remained a fundamental open question in episodic memory research. We addressed this by recording single neurons across two episodic paradigms that vary in task demands. Time cells and ramping cells were recruited in both paradigms, but with tuning properties that differed systematically: time fields were narrower in free recall, where temporal context must be internally generated, and ramping subtypes were asymmetrically distributed across paradigms in a pattern consistent with different demands on within-list (SR) versus within+across list (FR) temporal organization. Phase coding was more prominently engaged in serial reconstruction, organizing around both experimenter-defined item positions and empirically discovered population states, while free recall showed a graded phase relationship to population dynamics rather than to item positions. At retrieval, we observed evidence consistent with temporal context recovery supporting ecphory, as time cells led the broader assembly population within individual theta cycles across both paradigms, assemblies containing time cells reactivated preferentially and this difference scaled with recall success in free recall, and multi-neuron sequences initiated by time cells correlated with contextually organized behavior. Together, these findings reveal that temporal context representation and recovery in human medial temporal cortex recruit complementary cellular codes whose engagement depends on the structure the task provides.

A key novel aspect of our results pertains to neuronal sequences. Multi-neuron sequences that recurred across encoding and retrieval, were structured by assembly membership, and correlated with behavioral measures of contextually organized recall are the central novel finding of this work. Sequences have been well characterized in rodent hippocampus during navigation and sleep consolidation ^28,29,44^ and have recently been identified in human medial temporal cortex ^30^, but a direct link to the behavioral expression of episodic retrieval had not been established. Sequence frequency tracked temporal clustering in free recall and order accuracy in serial reconstruction, indicating that the same cellular mechanism serves contextually organized recall across paradigms. Within this framework, sequences initiated by a time cell were stronger predictors of recall organization than sequences broadly, both across and within subjects — the pattern the temporal context model predicts if temporal context must be recovered before item content can be retrieved ^1,2^, making sequences that begin with a context-carrying neuron the cellular instantiation of that recovery event. That these sequences are structured by assembly membership and engage time cells preferentially within their tuned time fields indicates that replay is organized by the population-level co-firing structure established at encoding ^22–24^.

The temporal context model has long predicted that retrieval begins with reinstatement of the contextual state present at encoding, which then re-cues the items bound to that context ^1,2^. Our data offer what is, to our knowledge, the first *direct cellular evidence* of this prediction: time cells fired at a systematically earlier theta phase than the surrounding assembly population during individual reactivation events, across both paradigms. The within-cycle architecture has a clear comparative precedent in rodent theta sequences, where place cells encoding the earliest-traversed locations occupy the leading phase of each cycle ^26,45–47^. In our paradigm, however, the leading-phase role is fixed to a functionally defined cell class (i.e. time cells) rather than rotating with the direction of spatial traversal ^48^, suggesting that the theta-sequence principle generalizes from spatial navigation to episodic retrieval in a form tied to the unidirectional temporal structure of episodic experience, with behavioral association to support its relevance to memory ^35,38^. That the same modulation was absent in serial reconstruction, where item identity is externally provided, is consistent with the interpretation that it is internally *reinstated temporal context*, rather than retrieval effort per se, that determines whether the time-cell-led assembly architecture is expressed.

Rate coding via context–sensitive neuron populations (time and ramping cells) and phase coding relative to theta oscillations have been treated as largely parallel accounts of temporal information in human MTL ^25,32,33^. Our data suggest they are complementary codes the recruitment of which depends on task demands. Phase locking was more prominently engaged in serial reconstruction than free recall, and exhibited item–specific phase coding at the population level. But neither paradigm produced the monotonic phase-position shift that classical theta-gamma accounts predict ^25,32^, a null consistent with Liebe et al. ^34^, who argued that apparent phase-serial position relationships reflect stimulus-timing interactions with oscillatory frequency rather than an active encoding mechanism. The use of HMM–defined states suggested that, when subjects must fully construct temporal context internally (i.e. in free recall), phase coding aligns to the brain’s discovered temporal groupings– a pattern consistent with chunking theories and indicating that phase-based coding depends on an internally structured temporal representation.

In the Curb (naturalistic) paradigm, ramping cells were identified at multiple timescales — spanning the full episode and resetting at scene boundaries. This heterogeneity is consistent with the Laplace-domain proposal that populations with different time constants collectively implement a continuous, scale-invariant representation of elapsed time ^18,49,50^. We found both upward and downward ramping subtypes were present at each timescale, with within-scene downward ramps the most prevalent. The aforementioned framework has largely been developed around rising ramps as a model of prospective temporal representation, but the functional role of declining ramps in human MTL remains poorly characterized and warrants direct investigation ^18,19^. The prevalence of ramping activity across session and scene timescales in the absence of structured retrieval demands suggests that this continuous temporal representation operates as a default property of the medial temporal system rather than being recruited specifically for episodic encoding — a view consistent with Cao et al. ^21^, who showed in rodent mPFC that anticipatory ramping requires an expected terminal event, implying that the ramping operates continuously while its prospective organization depends on behavioral demands.

Several aspects of our findings warrant caveats. We observe coordinated firing rather than a controlled perturbation, and the work does not establish causality. Our 25 ms-binned PCA-ICA assembly detection cannot exclude finer-grained co-firing structures with different phase relationships. We also lack a single-unit readout of item *content* against which the item-recovery side of context-driven retrieval can be tested directly. Concept-cell-style identification of individual items requires paradigms with extensive stimulus repetition, and our findings are complementary to such paradigms. Rey et al. ^51^ reported that most identity-responsive neurons in human MTL did not change firing across narrative contexts, while Bausch et al. ^52^ subsequently identified a small subpopulation with conjunctive context-by-stimulus tuning. Yuan et al. ^53^ recently argued that rodent hippocampal sequential activity beyond the first few seconds of a delay does not reliably support working memory, while Xiao et al. ^54^ document time-cell involvement in working-memory maintenance in humans under a distinct paradigm.

Overall, the identification of neuronal mechanisms supporting episodic memory creates opportunites to assess and design neuromodulation strategies. Specifically, the use of stimulation to induce phase may take advantage of the hidden structure in temporal tasks. However, a key outstanding question is whether these mechanisms extend to personally salient, natural autobiographical memories outside of lab environments. Detecting such data remains a key goal of ongoing research.

## Methods

### Participants

Single units were recorded from 62 patients with drug-resistant epilepsy undergoing invasive intracranial monitoring for seizure localization at one of two academic medical centers: Thomas Jefferson University Hospital (TJUH) or the University of Texas Southwestern Medical Center (UTSW). Institutional review board approval was obtained from both institutions, and written informed consent was provided by all participants. Electrode placement and number were determined solely by clinical criteria for seizure localization, independent of research considerations.

Two non-overlapping patient cohorts performed different memory tasks. Twenty-seven patients, recorded at both centers, performed the free-recall task. Thirty-five patients, recorded at UTSW, performed the serial-reconstruction task. The combined cohort was aged 39.0 ± 11.1 years (mean ± s.d.), with 17 female participants. A subset of the UTSW patients (n = 12) additionally completed a naturalistic viewing session (described in Behavioral tasks).

Following established procedures in our laboratory, each recording session was treated as an independent sample for statistical analysis ^15,36^.

### Electrophysiological recording

Patients were implanted with Behnke–Fried depth electrodes, which carry macroelectrode contacts along the shaft together with nine 40-*µ*m platinum–iridium microwires extending from the tip to resolve single-unit activity. The broadband microwire signal was digitally sampled at 30 kHz using a NeuroPort system (Blackrock Microsystems) at UTSW or at 32.6 kHz using a Cheetah system (Neuralynx) at TJUH. Microwire localization was performed by co-registering post-implantation computed tomography with pre-operative magnetic resonance imaging. Anatomical assignments were confirmed by a neuroradiologist using established anatomical landmarks of the medial temporal lobe, including the hippocampal head and body, collateral sulcus, and internal landmarks visible on MRI. All subsequent analyses were restricted to microwires located within the medial temporal lobe, which were classified into one of four subregions: the hippocampus proper, entorhinal cortex, amygdala, or parahippocampal gyrus. Hippocampal microwires were further subdivided into anterior and posterior groups, with the boundary determined at the level of the uncal notch. Electrode locations were visualized and rendered using BrainNet Viewer ^55^. Recording sessions containing clinically evident seizure activity or aura were excluded from analysis. Channels exhibiting excessive noise or frequent interictal spike discharges were discarded following clinical review.

### Spike detection and sorting

Before spike detection, the broadband signal was band-pass filtered from 30 kHz (or 32.6 kHz) to 1,000 Hz to isolate the high-frequency components containing single-unit activity, and volume-conduction subtraction was subsequently applied to attenuate the frequency-dependent noise shared across neighboring microwires ^15^. Spikes were then detected and clustered using Combinato ^56^. Every candidate unit was manually curated according to five quality criteria: (1) the shape and temporal stability of the mean spike waveform, (2) the proportion of inter-spike intervals shorter than 3 ms, serving as a measure of refractory violations, (3) the shape of the inter-spike-interval distribution, (4) the stationarity of the firing rate across the recording session, and (5) the degree of separation from other units on the same microwire. On the basis of these criteria, candidate clusters were either merged to represent a single neuron or discarded before any further analysis. To prevent duplicate counting of neurons detected across multiple microwires, unit pairs whose spike trains exhibited >50% coincidence (indicating temporal overlap exceeding physiological plausibility) were merged into a single unit. For microwires yielding multiple distinct units, pairwise isolation distances were computed to quantify the separation of clusters in spike-feature space. A final dataset of 768 units (free-recall cohort) and 247 units (serial-reconstruction cohort) was retained for analysis. The terms “units,” “cells,” and “neurons” are used interchangeably throughout to refer to individual single-unit recordings. For all units isolated, we computed units isolated per channel, mean firing rate, percentage of ISI less than 3 ms, modified coefficient of variation, signal-to-noise ratio of both the peak of the mean waveform and the mean of the mean waveform, and isolation distance distributions, which were all comparable with a reference high-quality dataset. These data are displayed in Supp. Figure 1.

### Behavioral tasks

In the free-recall task, patients studied lists of monosyllabic nouns presented one at a time on a bedside laptop and then recalled them aloud, a paradigm we have described previously ^15,36^. Each word remained on screen for 1.6 s, and a jittered interval of 0.8 to 1.2 s separated neighboring words. Lists contained 15 words at TJUH and 12 words at UTSW. After each list, patients completed an arithmetic distractor for at least 20 s, which discouraged rehearsal. All problems followed the format *A* + *B* + *C* =?. Patients then recalled as many words as possible, in any order, during a 30-s (UTSW) or 45-s (TJUH) retrieval period, and completed 4 to 25 lists per session.

In the serial-reconstruction task, patients studied lists of 10 monosyllabic nouns, with each word presented for 1.6 s and followed by a 1-s blank screen, and then performed a 30-s arithmetic distractor task. At test, all of the studied items reappeared together in a randomized array, and patients reconstructed the studied order by dragging into one of ten sequential boxes.. We scored an item as correctly recalled only when it occupied its correct absolute serial position within a run of at least two consecutive, correctly placed items. Patients completed 20 to 25 lists per session.

For the naturalistic viewing control, a separate group of UTSW patients watched a single episode of the television program *Curb Your Enthusiasm* lasting approximately 28 minutes. After viewing the episode, participants were presented with individual still images (72 total) and estimated when each occurred on a linear scale.

### Local field potential processing

For analyses requiring spectral phase or power, the microwire signal was low-pass filtered at 300 Hz using a second-order Butterworth filter, effectively removing spike-related artifacts while preserving local field potential. A second-order Butterworth notch filter was subsequently applied at 60 Hz to attenuate line noise. The signal was then downsampled to 1 kHz to reduce computational demands while maintaining sufficient temporal resolution for phase estimation.

Instantaneous phase and power were extracted by convolving the filtered signal with six-cycle Morlet wavelets centered at analysis-specific frequencies, a wavelet-based approach chosen to optimize time-frequency resolution for examining neural oscillations ^15,36^.

### Time cell identification

Time cells were identified using a previously published nonparametric procedure optimized for detecting temporally tuned neurons in human medial temporal lobe recordings ^15^, following established approaches in the time-cell literature ^11,12^. The procedure tested whether each neuron’s firing rate varied significantly with elapsed time within the encoding interval without assuming a particular form for the temporal tuning relationship. For each neuron, the session-wide spike train was convolved with a Gaussian kernel (standard deviation: 0.5 s) to obtain a continuous estimate of instantaneous firing rate, referred to as the tuning curve. For each encoding list, the corresponding segment of the tuning curve was divided into contiguous, non-overlapping time bins (bin width: approximately 1 s), and the mean firing rate was computed within each bin, yielding a list-by-bin matrix. Temporal modulation across bins was tested using a Kruskal–Wallis test, a rank-based nonparametric test robust to the non-normal, Poisson-distributed nature of neural spike counts. Neurons meeting the threshold of p < 0.05 were advanced to a permutation test to determine whether their temporal tuning exceeded chance levels. For each neuron, a null distribution of test statistics was generated by circularly shifting the session-wide tuning curve by a randomly selected amount (1,000 shuffles), then re-binning and re-testing. Circular shifting preserves the temporal structure and autocorrelation of the neuron’s firing while disrupting the alignment of spikes to the behavioral epoch, isolating genuine time-dependent firing. A neuron was classified as a time cell when fewer than 5% of shuffles yielded a smaller *p*-value than the unshuffled data. This identification procedure was applied identically to both memory-task datasets and, for retrieval analyses, to tuning-curve segments spanning each retrieval period. Each time cell was further classified by the behavioral context in which it achieved significance: encoding only, retrieval only, or both periods.

Time fields were localized within each identified time cell’s trial-averaged tuning curve for analyses requiring specification of a temporal receptive field, including cross-task comparisons of field widths. The tuning curve was re-binned at finer temporal resolution (bin width: approximately 0.2 s), and contiguous runs of consecutive bins were identified whose mean firing rate exceeded the neuron’s session-wide mean firing rate. A run was designated as a time field when two conditions were met: (1) the run spanned a minimum contiguous duration of 1 s, and (2) the neuron exhibited firing within that time window on at least half of the lists within that task. Because time fields were defined exclusively for neurons already classified as time cells, cross-task comparisons of field widths were made between the independently identified free-recall and serial-reconstruction populations, ensuring that any task-dependent differences in field properties reflected genuine variations in temporal tuning rather than differences in the composition of identified time cells.

### Ramping cell identification

Ramping cells were identified using an established approach that has been applied across diverse species and brain regions, including rodent hippocampus, non-human primate prefrontal and parietal cortex, and human medial temporal and prefrontal cortex ^18–20^. The procedure tests whether a neuron’s firing rate exhibits monotonic changes over extended timescales, either across the entire encoding epoch or the full recording session, independently of trial-to-trial stimulus fluctuations. For each neuron, spikes were counted in non-overlapping 1-second bins spanning the entire session. The resulting spike counts were modeled using a Poisson generalized linear model (GLM) with log link. The model included two primary predictors of interest capturing different temporal scales: (1) session time, measured continuously from recording onset, and (2) epoch time (or scene time in the naturalistic viewing task), measured within each encoding unit. To account for task structure and event-related firing, the model additionally included nuisance regressors: epoch (or scene) identity, word onset, stimulus presentation or word visibility (for free-recall and serial-reconstruction tasks), list (or scene) onset, and—in free-recall only—retrieval onset and vocalization onset. All continuous predictors were standardized (z-scored) prior to fitting. Model selection was performed using stepwise regression (entry threshold: p < 0.01; removal threshold: p < 0.005). A neuron was classified as a ramping cell when two criteria were satisfied: (1) the final selected model retained at least one temporal predictor (session or epoch time), and (2) that predictor’s contribution was validated against a null distribution generated by 1,000 circular shifts of the spike train. Circular shifting preserves spike-train autocorrelation and temporal structure while breaking the true alignment between spikes and elapsed time, providing a robust test of whether temporal modulation exceeds what would arise from autocorrelated noise.

Each identified ramping cell was labeled by the direction and timescale of its temporal coefficient. Cells were designated as session-ramping-up or session-ramping-down when the session-time coefficient indicated increasing or decreasing firing across the full recording, respectively. Similarly, cells were designated as epoch-ramping-up or epoch-ramping-down when the epoch-time coefficient indicated increasing or decreasing firing within each encoding epoch, respectively. When both temporal predictors were retained in the final model, the cell was labeled based on the predictor with the larger absolute coefficient, indicating the dominant timescale of temporal organization. For visualization, the model’s predicted firing rate (derived from all retained predictors) was overlaid on the observed firing rate across the session.

### Population decoding of elapsed time

To determine whether elapsed time within the encoding interval was decodable from population firing rates, a multinomial logistic-regression decoder was trained to predict the time bin from population activity vectors. Each encoding list was divided into 20 equal temporal bins. For each bin, a population firing-rate vector was constructed containing the spike rate (spikes per second) of all simultaneously recorded (or pseudo-simultaneously recorded) neurons. Each vector underwent sequential preprocessing: square-root transformation to stabilize variance; z-score standardization within each neuron to remove baseline firing-rate differences; and temporal smoothing across adjacent bins using a Gaussian kernel (standard deviation: 0.5 bins) to reduce noise while preserving temporal structure. The decoder was implemented as a multinomial logistic-regression classifier using scikit-learn ^57^.

Because neurons were recorded across separate sessions rather than simultaneously, pseudo-populations were constructed to test population-level temporal coding. Within each cross-validation fold, 200 pseudo-trials were generated by independently sampling, for each neuron, a firing-rate vector from a randomly selected list. This approach permitted assessment of population decoding while preserving the statistical structure of neural responses within individual encoding lists. The classifier was evaluated using tenfold cross-validation stratified by list, ensuring that no list contributed data to both the training and test sets within a given fold. Chance performance was estimated by repeating the full cross-validated decoding procedure 100 times with time-bin labels randomly shuffled, generating a null distribution of classifier accuracy. The decoder was run independently on four distinct neuronal populations: time cells, ramping cells, the union of time and ramping cells, and cells belonging to neither category. This stratification was performed separately for each task (free-recall and serial-reconstruction) and on the combined sample. Performance was quantified using two complementary metrics: (1) classification accuracy (percentage of correct bin assignments), and (2) mean absolute error between predicted and true bin.

To determine the temporal localization of decodable time information, decoding accuracy was compared between the first (bins 1–10) and second (bins 11–20) halves of the encoding interval with a one-sided Mann–Whitney U test across cross-validation folds. To characterize the structure of decoding errors, accuracy was scored at progressively coarser temporal tolerances: (1) exact bin prediction, (2) prediction within ±1 bin of the true bin, (3) prediction within ±2 bins, and (4) prediction within the same temporal third of the list (primacy: first third; middle: second third; recency: final third). Because the free-recall neuronal population substantially exceeded the serial-reconstruction population in size, free-recall decoding was repeated after subsampling neurons to match the serial-reconstruction population size. For this subsampling analysis, 100 independent random samples were drawn, with neurons retained in proportion to each session’s contribution to the original population, ensuring that session-level effects were preserved across resamples.

### Cell assembly identification

Cell assemblies were identified following previously established procedures for human medial temporal lobe recordings ^36^, adapted from a principal- and independent-component analysis framework developed in rodent studies ^37^. Analysis was restricted to neurons with baseline firing rates *≥* 0.5*Hz*. For each recording session, spike trains were binned at 25-millisecond resolution during encoding (a timescale chosen to match the gamma band), and z-scored to prevent high-firing rate neurons from dominating the analysis. The resulting neuron-by-bin matrix was subjected to principal component analysis with components retained if their eigenvalues exceeded the upper bound predicted for independent firing by the Marchenko–Pastur distribution, thereby isolating genuine co-firing structure. The firing-rate matrix was then projected onto retained components and independent component analysis (fastICA) was applied, so that the number of recovered assemblies equaled the number of significant components. Each assembly was described by a weight vector, normalized to unit length and signed such that its maximum weight was positive. Neurons were classified as assembly members if their absolute weights exceeded the ensemble mean plus one standard deviation. Assemblies were required to contain at least two member neurons to exclude single-neuron contributions. To validate that the detected assemblies reflected genuine neural organization rather than thresholding artifacts, the observed number was compared against the null distribution obtained from circularly shifted spike trains.

The moment-to-moment expression of each assembly was quantified as as

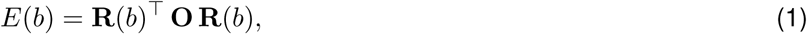

where **R**(*b*) is the vector of z-scored firing rates in time bin *b*, and **O** is the outer product of the assembly’s weight vector with its diagonal set to zero, which ensures that no single neuron’s firing alone drives the measure.

### Assembly reactivation

Assembly reactivation events during retrieval were defined as the time bins in which an assembly’s expression strength exceeded its session-wide mean by more than two standard deviations ^38^. To test whether assemblies carrying temporal context were preferentially reactivated, the reactivation strength of assemblies that contained at least one time cell was compared to that of assemblies that contained none. To determine whether the reactivation strength difference depended on memory success, a median split was used to divide each session’s lists into low- and high-recall halves, and a linear mixed-effects model was fit to determine whether the reactivation-strength difference between time-cell-containing and time-cell-lacking assemblies varied with recall level. A by-subject random intercept was included in the model to account for the multiple assemblies contributed by each patient.

### Time-cell theta phase at assembly reactivation

To determine when time cells fired relative to the assemblies they accompanied, we restricted our analysis to time cells that were not members of the assembly. We then examined the theta phase of these non-member time-cell spikes around each reactivation event. For each assembly, a single preferred theta frequency was identified by computing each member neuron’s phase-locking strength (mean resultant length) at every frequency from 2–10 Hz, averaging these values across the assembly’s members, and selecting the frequency that maximized the assembly-averaged phase-locking. This yielded one preferred frequency per assembly rather than per neuron. For each reactivation event, the instantaneous theta phase was extracted for each time-cell spike occurring within ±150 ms of the event, and the circular mean of these phases was compared to the mean theta phase of the assembly’s constituent neurons. Negative phase differences indicated that time cells fired at earlier phases of the theta cycle relative to assembly neurons. To avoid circular analysis, time-cell phases were drawn exclusively from neurons that were not members of the assembly under consideration The phase-locking relationship for each assembly was characterized by the circular mean and resultant vector length of its event-wise phase differences. Consistency of this relationship across assemblies was quantified with the mean resultant length; a Rayleigh test evaluated whether the phase distribution differed significantly from uniform. Statistical significance was assessed with a permutation test that resampled subjects with replacement (10,000 iterations) to preserve the hierarchical structure of assemblies within patients, using the circular-statistics toolbox. The identical analysis was applied to ramping cells, testing their theta-phase relationship to assembly reactivation. As a robustness check, the time-cell lead was recomputed at fixed theta frequencies from 3 to 9H̃ z, across a range of activation-window widths, and directly in spike time without assuming an oscillation (Supp.F̃igureS̃ 3C–F).

### Within-assembly gamma phase

To determine whether temporal structure within assemblies was evident at faster timescales, gamma-band phase relationships between time-cell and non-time-cell members of assemblies were examined ^36^. Gamma phase was extracted using six-cycle Morlet wavelets centered at 40 Hz (period: 25 ms, matched to the assembly detection timescale to assess phase relationships at equivalent temporal resolution). For each assembly, each member neuron was summarized by the circular mean of its gamma phases (pooled across all of its spikes in the encoding and retrieval windows). Within each assembly, the circular mean of all pairwise phase differences between time-cell and non-time-cell members was then computed (negative = time cells lead), yielding one value per assembly, separately for low- and high-recall lists. Across assemblies, the per-assembly phase differences were summarized by their circular mean and mean resultant length, with non-uniformity assessed by a Rayleigh test. Significance of the time-cell-versus-non-time-cell difference was further evaluated with a subject-resampling bootstrap (10,000 iterations) and a permutation test (5,000 permutations)that shuffled the time-cell and non-time-cell labels of members within each assembly. Between-condition differences in mean phase were compared using the Watson–Williams test, a parametric circular statistics test applied only when mean resultant length exceeded 0.45, indicating sufficient phase concentration for the test’s distributional assumptions.

### Spike sequence detection

Repeating spatiotemporal spike sequences were detected using SPADE, an established framework that identifies neuronal firing patterns occurring more frequently than expected by chance based on spike-train properties ^40,41,58^. Each recording session was analyzed on a unified timeline that concatenated the encoding and retrieval periods of all lists, permitting the detection of sequences that recurred across both periods as a single pattern. Spikes were binned at 5 ms resolution, and patterns spanning up to 300 ms and involving at least three neurons were identified. Each candidate pattern was evaluated against a null distribution generated by dithering spike times (1,000 surrogates with jitter drawn from a Poisson process to preserve spike-train statistics while breaking temporal structure). Multiple comparisons across the pattern spectrum were corrected using the Benjamini–Hochberg false discovery rate procedure ^59^ at *α* = 0.05. Because the number of candidate patterns grows steeply with the number of recorded spikes, he minimum spike count and minimum number of occurrences per candidate pattern were set on a per-session basis, using the most permissive thresholds that kept the candidate set computationally tractable. These settings governed only which patterns were submitted to surrogate testing. Final significance was determined by the dither-based surrogate test itself. Specifically, each pattern’s signature was compared against the surrogate distribution to obtain a p-value, corrected across the pattern spectrum with the Benjamini–Hochberg FDR procedure (*α* = 0.05), and followed by pattern set reduction to remove redundant sub- and super-patterns. A sequence that recurred in both the encoding and retrieval periods of the same list was designated a replay event, the cellular signature through which an encoded sequence can be reinstated at retrieval.

Significant sequences were characterized by their size (number of participating neurons), duration, and frequency of occurrence. For each subject, the median value of each property was calculated separately for each task, and the two tasks were compared using Mann–Whitney U tests. To determine whether time cells were engaged at their tuned moments when participating in sequences, the fraction of sequences in which a time-cell participant fired within its established time field was measured. This observed fraction was compared to the expected fraction under the null hypothesis that sequence participation and time-field firing were independent, using a Wilcoxon signed-rank test across subjects.

### Assembly and sequence relationships

For each subject, a Fisher’s exact test was used to compare the proportion of sequence-participating neurons that were assembly members with the proportion of assembly members in the recorded population, determining whether sequences preferentially recruited neurons with assembly membership (Supp.F̃igureS̃ 4A).

The degree to which sequence-participating neurons were drawn from the same assembly was assessed by computing the assembly homogeneity for each sequence, quantified as the proportion of participating neurons belonging to the dominant assembly divided by the total number of participating neurons. The observed homogeneity was compared to a null distribution generated by randomly shuffling assembly-membership labels—reassigning each sequence-participating neuron to a randomly selected assembly while preserving assembly sizes and identities—testing whether temporal sequences exhibited non-random assembly structure (Supp.F̃igureS̃ 4B).

Finally, the proportion of sequences that recurred within a study list, meaning they appeared in both the encoding and retrieval periods of the same list, was compared between sequences containing at least one assembly member and sequences lacking assembly members using a two-sided Fisher’s exact test.

### Relating sequences to recall behavior

To determine whether the temporal organization of recalled items was related to detected sequences, the temporal clustering factor was computed—an established behavioral measure quantifying the tendency of successive recalls to originate from nearby encoding positions ^4,5^. For each free-recall list, the temporal clustering factor and the sequence recurrence rate were computed. A within-subject Pearson correlation was calculated between these measures after mean-centering both variables within each subject to remove between-subject variability. Significance was assessed with a subject-resampling bootstrap (5,000 iterations, resampling with replacement) that preserved the hierarchical structure of lists nested within patients.

The analysis was then repeated with sequence recurrence restricted to time-cell-led sequences, defined as sequences whose earliest-firing neuron was a time cell, testing whether temporal structure specifically driven by time cells predicted behavioral temporal clustering. For the serial reconstruction task, where temporal clustering is undefined, the fraction of items correctly reconstructed in their original encoding order was used as the behavioral measure. This was related to the fraction of detected sequences containing at least one time cell using an identical within-subject correlation and subject-resampling procedure. For visualization, lists were divided at each subject’s median of the relevant neural measure (replay rate; or, for the time-cell-specific analyses, the time-cell-led or time-cell-containing replay fraction), and the behavioral measure was averaged within each half. The corresponding behavioral measure was compared between the resulting upper and lower halves using paired t-tests or Wilcoxon signed-rank tests as appropriate.

### Theta phase locking

To determine whether non-temporal neurons exhibited consistent spike-phase relationships during encoding, a second-order phase-locking measure was computed for each neuron at each serial position (or hidden Markov model state). Analysis was restricted to non-temporal neurons (defined as those classified as neither time cells nor ramping cells) because time and ramping cells express phase precession, a distinct phase-related phenomenon that would confound assessment of spike-phase stability across positions.

For each neuron, the local field potential (LFP) recorded from the same microwire was decomposed into instantaneous power and phase at each frequency using six-cycle Morlet wavelets. For each frequency band of interest (delta, 2–5 Hz; theta, 5–9 Hz), a composite instantaneous phase was derived at each timepoint as the angle of the amplitude-weighted mean complex phasor across frequencies within the band:

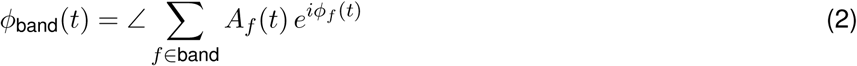

where *A_f_* (*t*) and *ϕ_f_* (*t*) denote the instantaneous amplitude and phase at frequency *f*, respectively. Within each encoding list, the LFP phase was assigned to every spike at each serial position at the moment of spike occurrence. A first-order mean resultant vector was computed across all spikes at each serial position within each list. Phase non-uniformity was tested using a Rayleigh test, which evaluates whether spike phases differ significantly from a uniform distribution (null hypothesis: phases are uniformly distributed around the circle).

The per-list mean phase angles were treated as independent observations. The mean resultant length (MRL) was computed across all lists to quantify the consistency of phase preference. Per-list Rayleigh p-values were combined across lists using Fisher’s method, yielding a single composite p-value. Neurons were required to contribute spikes in at least two lists to ensure sufficient statistical power for phase-locking assessment. A null distribution was generated for each neuron by circularly shifting each list’s spike train independently by a random amount (1,000 permutations). Circular shifting preserves the spike train’s autocorrelation and temporal structure while breaking its alignment to LFP phase, providing a conservative test of phase-locking that controls for spurious phase relationships arising from firing-rate fluctuations alone. A neuron was classified as phase-locked at a serial position when its observed MRL exceeded the 95th percentile of its permutation null (equivalent to p < 0.05). Population-level significance was assessed by comparing the observed count of phase-locked neurons against the distribution of null counts from the same shuffles. The population p-value was defined as the proportion of permutations in which the null count equaled or exceeded the observed count.

To determine whether preferred phase differed systematically across serial positions, an omnibus test was applied to the mean preferred phases of significantly phase-locked neurons at each position. For each serial position, the circular mean of preferred phases across all phase-locked neurons at that position was computed. A weighted circular between-group variance statistic was then calculated as

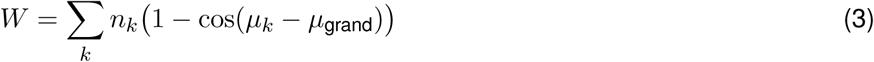

where *n_k_* is the number of phase-locked neurons at position *k*, *µ_k_* is their mean preferred phase, and *µ*_grand_ is the grand circular mean across all positions. Significance was assessed by randomly permuting all phase observations across positions (preserving position group sizes; 10,000 permutations) and comparing the observed *W* against the resulting null distribution. If the omnibus test was significant, pairwise permutation tests on the absolute angular difference between each pair of positions were conducted (10,000 permutations each), with Bonferroni correction for multiple comparisons.

The overlap between phase-locked neuronal populations at different serial positions was quantified using the Jaccard index, computed for each pair of positions as the size of the intersection divided by the size of the union of the two phase-locked neuron sets. Each observed Jaccard index was compared against a null distribution generated by randomly permuting each neuron’s phase-locking status across positions (10,000 permutations), preserving the number of phase-locked neurons at each position. Multiple comparisons across all position pairs were corrected using Bonferroni correction at *α* = 0.05.

### Circular-linear correlation

To test whether spike phase varied systematically with serial position, a circular-linear correlation was computed between instantaneous spike phase and serial position (or hidden Markov model state) for each neuron. Rather than analyzing serial positions independently, all spikes were pooled across serial positions, encoding lists, and tasks to maximize statistical power. Each spike was assigned two values: (1) its instantaneous LFP phase extracted from the analytic signal, and (2) the serial position (1–10 for serial reconstruction; 1–15 for free recall) or HMM state (1–4 for serial reconstruction; 1–3 for free recall) of the word during whose encoding window the spike occurred. The circular-linear correlation coefficient r was computed following established methods ^60^, defined as:

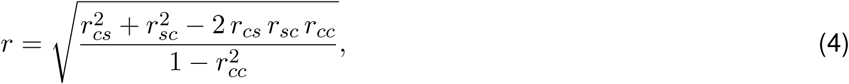

where *r_cs_* is the Pearson correlation between cos(*θ*) and serial position (or hidden Markov model state), *r_sc_* the correlation between sin(*θ*) and serial position (or hidden Markov model state), and *r_cc_* the correlation between cos(*θ*) and sin(*θ*). The coefficient ranges from 0 to 1, with larger values indicating a stronger monotonic phase-position relationship. Significance of each neuron’s phase-position relationship was assessed using the same circular-shift permutation procedure described above (1,000 permutations per neuron, preserving spike-train autocorrelation while disrupting phase-position alignment). A neuron was classified as exhibiting significant phase-position coupling when its observed r exceeded the 95th percentile of its null distribution (equivalent to p < 0.05, two-tailed). Population-level significance was determined by comparing the observed proportion of neurons meeting the significance threshold against the null proportion computed from the same permutations.

For both phase-locking and phase-position analyses, neurons were required to fire at least one spike in each analyzed position-and-list combination to ensure sufficient data for reliable phase estimation. Results are reported separately for the delta (2–5 Hz) and theta (5–9 Hz) frequency bands. A neuron was considered significant if it met the significance threshold in either band, reflecting the biological expectation that phase-serial-position relationships may be expressed at either timescale.

### Hidden Markov model

Neural population dynamics during encoding were modeled using a left-to-right event-segmentation hidden Markov model (HMM), following the framework of Baldassano et al. ^61^. The model was applied separately to the free-recall (FR) and serial-recall (SR) datasets, with single units pooled across subjects to form a pseudo-population for each task. Each encoding list constituted one trial. Spike trains were convolved with a normalized Gaussian kernel (*σ* = 250 ms) to estimate instantaneous firing rate. The smoothed firing rate trace for each neuron on each list was resampled to 530 uniformly spaced time points, after which 15 bins were trimmed from each edge to mitigate boundary artifacts, yielding 500 time points per trial. The firing rate of each neuron was then z-scored within each trial independently; neurons with zero variance on a given trial were excluded from the emission computation for that trial. Neurons missing data on 10 or more lists, and lists in which no neuron had data, were excluded prior to model fitting.

The HMM assumed a left-to-right state topology, whereby transitions were permitted only to the current state or the next state, enforcing a monotonically progressing sequence of K discrete neural states. Emission distributions were modeled as independent Gaussians over z-scored firing rates. Parameters were estimated via Expectation-Maximization (EM) with an initial self-transition probability of 0.95, an emission variance floor of 0.05, and a minimum advance probability of 0.003. Convergence was assessed using a tolerance of 1*x*10^−4^. The number of EM iterations was set to a maximum of 300 for FR and 200 for SR. Models were fit across a range of K = 3–11 states. For each value of K, 3-fold cross-validation was performed; the model was trained on two folds and evaluated on the held-out fold, with performance quantified as log-likelihood per time point on held-out data. The optimal K was selected as that which maximized cross-validated log-likelihood per time point. The Bayesian Information Criterion (BIC), computed on training data, was evaluated as a secondary check.

### Naturalistic viewing analyses

To determine whether temporally tuned neurons emerge outside the bounded, repeating structure of the memory tasks, single units were analyzed during a naturalistic viewing session in which patients watched a single episode of *Curb Your Enthusiasm*. To relate the episode’s temporal structure to behavioral timing, patients’ periodic judgments of elapsed time were examined. Each judgment was correlated with the true elapsed time using Pearson correlation, and the strength of this relationship was summarized across scenes with bootstrap confidence intervals (10,000 iterations), quantifying the fidelity of temporal perception during naturalistic viewing.

For ramping cells, the generalized linear mixed-effects model was adapted to the continuous temporal structure of the naturalistic episode. For each neuron, spikes were counted in one-second windows across the full episode. A Poisson generalized linear mixed-effects model was fit with predictors including elapsed time from the episode start, elapsed time within the current scene, and scene onset; a random intercept for scene captured baseline firing-rate differences between scenes. A neuron was classified as a ramping cell when the full model significantly improved over a reduced model containing only the scene random intercept (likelihood-ratio test, *α* = 0.05) and when at least one temporal predictor (episode time or scene time) contributed significantly (p < 0.05).

