## Supplemental Figures for "Time cells lead neural reinstatement of episodic memory"

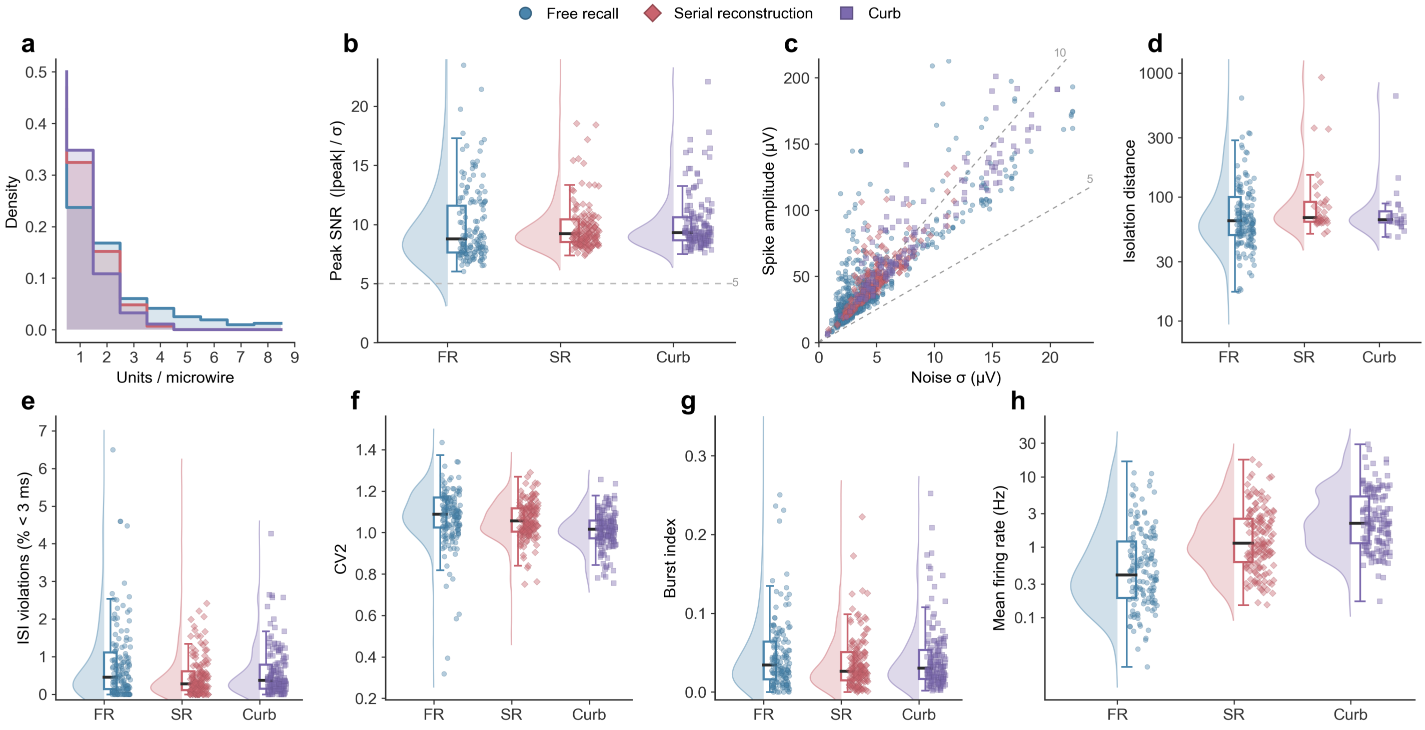


**Figure S1. Single-unit recording quality is equivalent across the three recording cohorts.** (**a**) Well-isolated units per microwire in the free-recall (FR), serial-reconstruction (SR), and naturalistic-viewing (Curb) cohorts. (**b**) Peak signal-to-noise ratio ($|\mathrm{peak}|/\sigma_{\mathrm{noise}}$); dashed line, the SNR $\geq 5$ inclusion criterion. (**c**) Spike amplitude versus background-noise $\sigma$; dashed rays, SNR ratios of 5 and 10. (**d**) Isolation distance, computed on multi-unit wires. (**e**) Inter-spike-interval violations (spikes $< 3$ ms apart). (**f**) Spike-train regularity ($\mathrm{CV}_2$). (**g**) Burst index. (**h**) Mean firing rate (log scale). Panels **b**–**h**, raincloud plots (box, median and IQR); cohort statistics, Table S2.

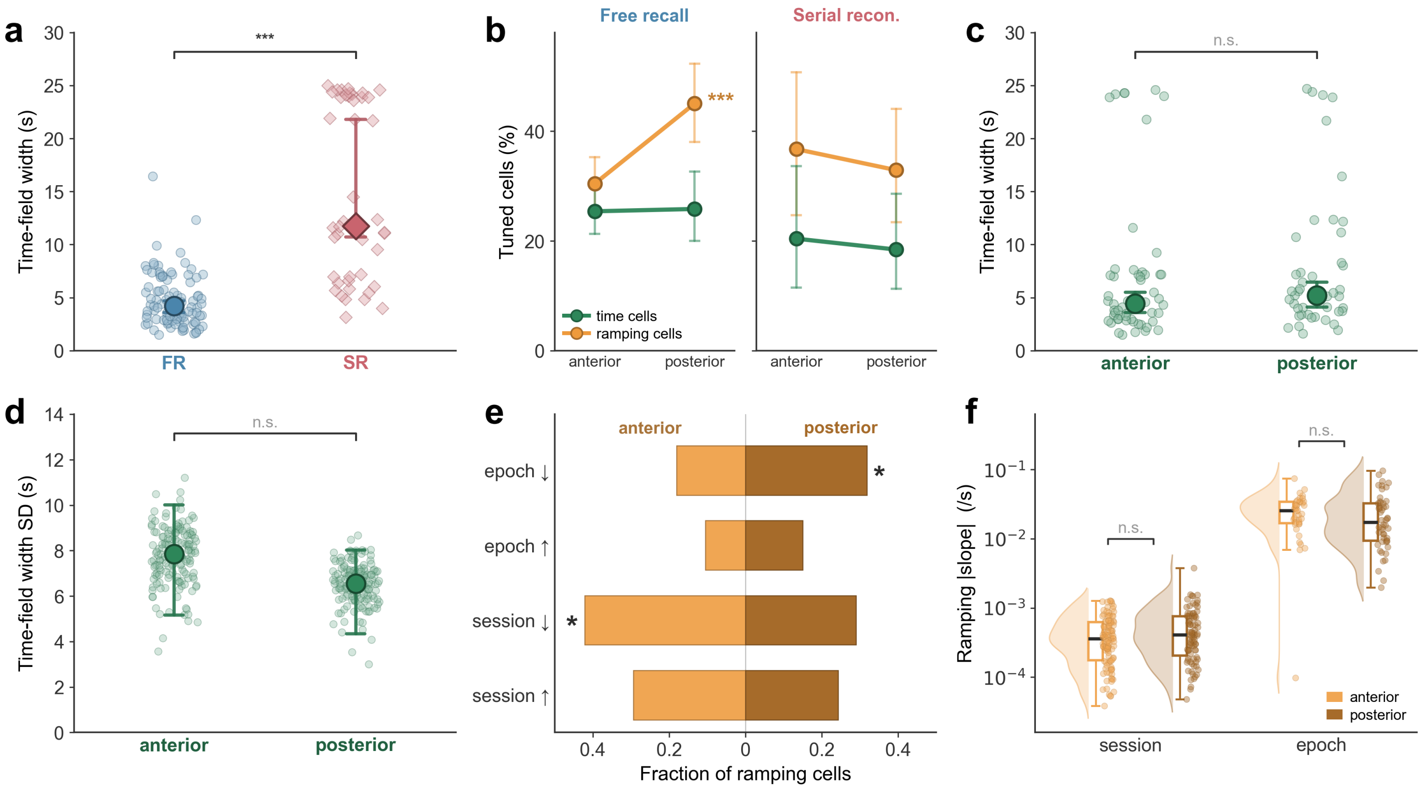


**Figure S2. Robustness of time-cell and ramping-cell tuning and its distribution along the hippocampal long axis.** (**a**) Mean time-field width in absolute time (seconds) for free recall (FR) versus serial reconstruction (SR). (**b**) Prevalence of time cells and ramping cells in anterior versus posterior hippocampus for FR (filled, solid) and SR (open, dashed), with 95% CIs. (**c**) Median and (**d**) standard deviation of time-field width, anterior versus posterior (FR and SR time cells pooled; n.s.). (**e**) Ramping-cell counts by direction (up/down) and timescale (session/epoch), anterior versus posterior. (**f**) Ramping-slope magnitude (absolute stepwise-GLM time coefficient, log scale) for session- and epoch-time ramping, anterior versus posterior (n.s.).

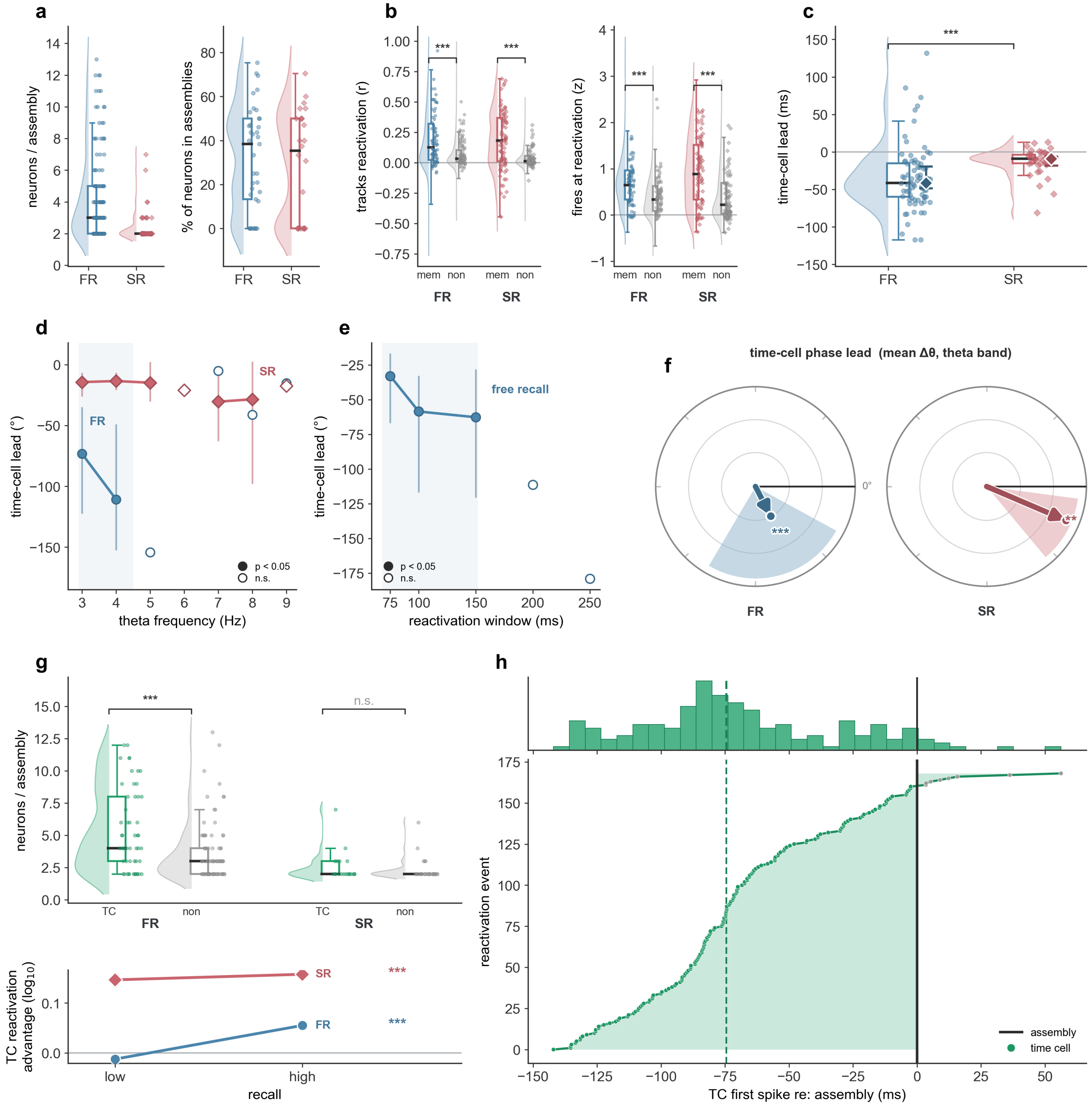


**Figure S3. The time-cell phase lead in assembly reactivation is robust and not explained by assembly size or firing rate.** (**a**) Member neurons per assembly and the percentage of recorded neurons in at least one assembly, per session, in free recall (FR) and serial reconstruction (SR). (**b**) Per unit, correlation with its assembly's reactivation time course (left) and firing at its reactivation peaks (right), for assembly members versus non-members. (**c**) Time-cell lead measured directly in time, no oscillation assumed: how far the time cell's first spike precedes the rest of the assembly (negative, time cell earlier). (**d**) The same lead as theta phase ($\Delta\theta$), recomputed at fixed theta frequencies from 3 to 9 Hz (filled markers, statistically resolved). (**e**) The same lead across activation-window widths in free recall (150 ms window of Figure **??** marked). (**f**) Mean theta-phase lead per paradigm (compass plot). (**g**) Top, assembly size with versus without a time cell; bottom, time-cell reactivation-strength advantage at low versus high recall. (**h**) Reactivation events of one example assembly, each showing the time cell's first spike relative to the assembly (line at 0).

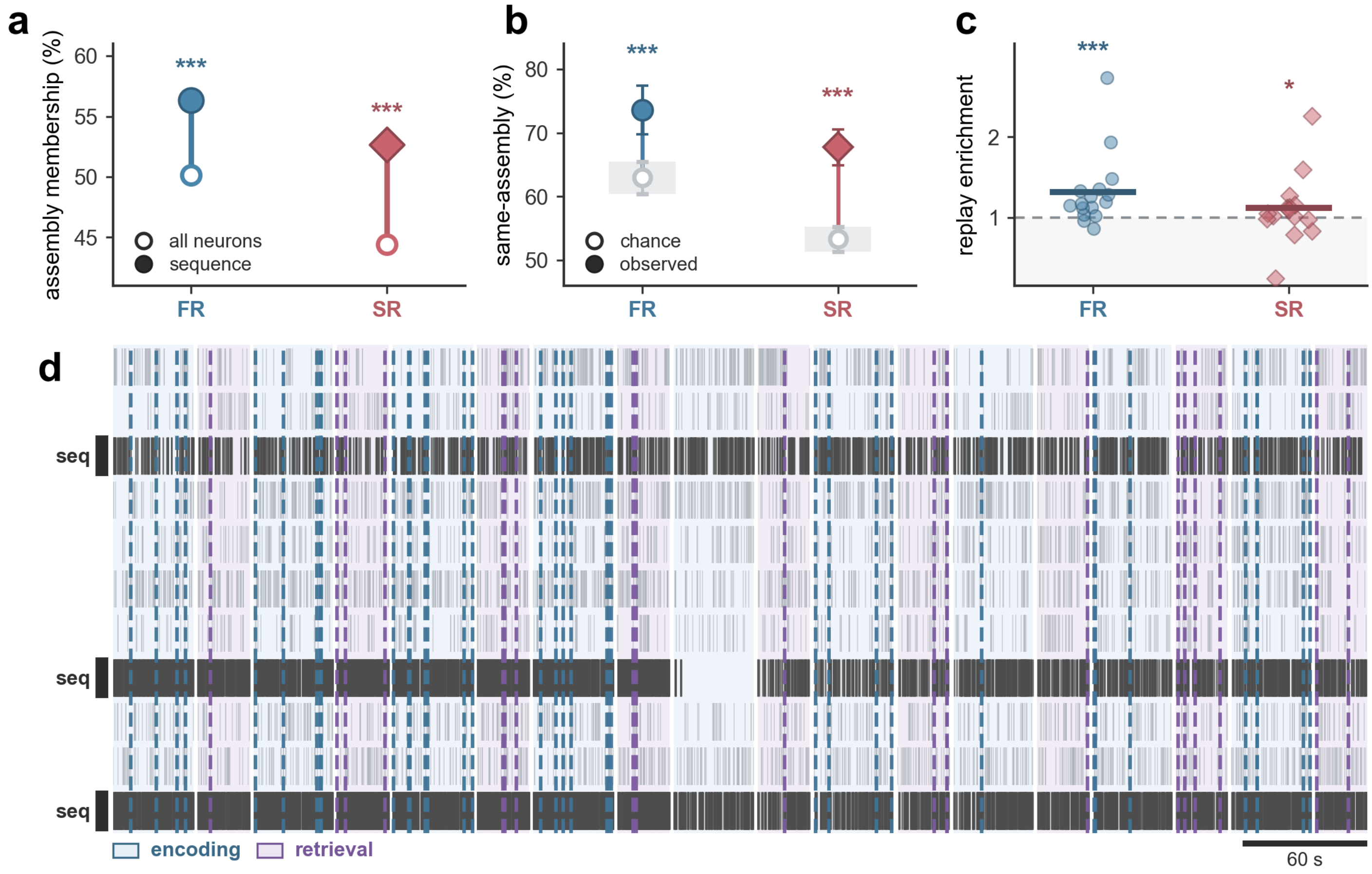


**Figure S4. Multi-neuron sequences are built from the encoding assembly architecture and recur above chance.** (**a**) Assembly membership of all recorded neurons (open) versus sequence-participating neurons (filled), in free recall (FR) and serial reconstruction (SR). (**b**) Within-sequence homogeneity: how often the neurons of a sequence belong to the same assembly (filled) versus a label-shuffle null (open). (**c**) Per-subject replay enrichment: the rate at which a sequence recurs in both encoding and retrieval of a list, relative to a dither-surrogate null (dashed line, chance). (**d**) Example replay raster (subject TJ042_2): one recurring three-neuron sequence (dark, "seq") among other simultaneously recorded neurons (gray) across the encoding (blue) and retrieval (purple) periods of nine consecutive lists; dashed lines, sequence occurrences.

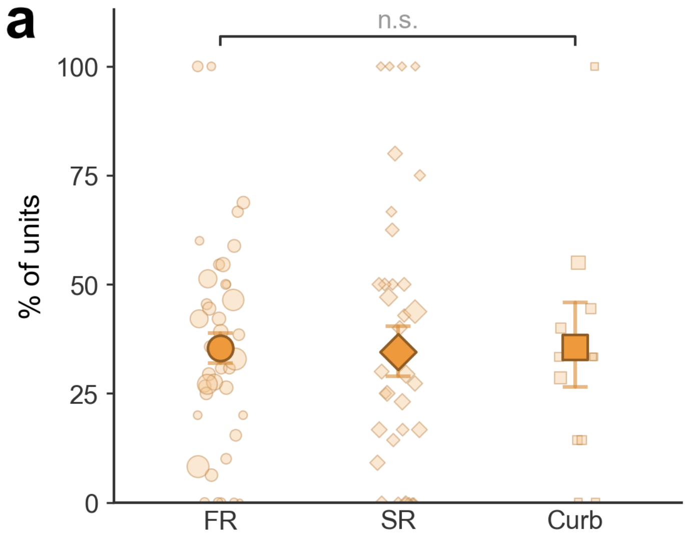


**Figure S5. Ramping cells are recruited at comparable rates across paradigms.** Proportion of recorded units classified as ramping cells in free recall (FR), serial reconstruction (SR), and naturalistic viewing (Curb). Faint markers, individual sessions (size proportional to units per session); bold markers and whiskers, pooled proportion and 95% CI (n.s.).
